# Phenolic compounds in *Medicago truncatula* roots are under the influence of *Agrobacterium fabrum* through its species specific-genes regions

**DOI:** 10.1101/2024.12.27.630484

**Authors:** Rosa Padilla, Ludovic Vial, Florence Dechèvre, Nguyen Thi Huyen Thu, Vincent Gaillard, Guillaume Meiffren, Gilles Comte, Xavier Nesme, Céline Lavire, Isabelle Kerzaon

## Abstract

The impact of plant microbiota on the health and physiology of their host is increasingly studied and recognized. However, in the rhizosphere, the functions of most bacteria and the genetic determinants involved in the molecular dialogue between plant and bacteria are poorly understood. Agrobacteria are ubiquitous soil borne and rhizospheric bacteria able to establish commensal or even beneficial interactions with plant roots. The genomic species *Agrobacterium fabrum* harbor seven specific-regions (SpG8-1 to SpG8-7), whose annotation seems to indicate a close connection with the plant during plant-bacteria interactions. To evaluate the involvement of *A. fabrum*-specific regions in the plant-bacteria interaction, deletion mutant strains of each *A. fabrum*-specific region were inoculated on *Medicago truncatula* roots. Root metabolite profiles were compared by UHPLC-UV/DAD-ESI-MS QTOF analyses, and the highlighted discriminating metabolites were annotated by tandem mass spectrometry. Metabolomic analyses have shown that *A. fabrum* inoculation modulates the content of phenolic compounds in *M. truncatula* roots, in particular flavonoids. These root metabolite modulations observed with the wild-type strain often appear to be linked to at least one of the *A. fabrum*-specific genes, as almost all *A. fabrum*-specific regions showed an influence on one or more of these specialized root metabolites. In addition, our results underlined a putative cross-talk or coordinated effect of the *A. fabrum*-specific regions during the interaction of *A. fabrum* with *M. truncatula*, as all mutants except one induced similar modifications on flavonoids. These results contribute to a better understanding of the ecological niche construction of *A. fabrum* highlighting the importance of its specific genes in the establishment of this fine-tuned interaction.

## INTRODUCTION

The critical role and diversity of the plant microbiota have gained widespread recognition. Microorganisms associated with the plant can profoundly impact their health: ranging from causing devastating diseases to promoting growth, health, and productivity. For certain well-studied bacteria, such as *Xanthomonas* (a notorious plant pathogen) or nitrogen-fixing rhizobia, we now have a solid understanding of their effects on plants (Philippot et al., 2013). For model bacteria or bacteria of economic interest, key genetic determinants driving these interactions have been identified, shedding light on the mechanisms underlying their influence. In the rhizosphere, however, the functions of most bacteria and the genetic determinants involved in the molecular dialogue between bacteria and plants are poorly understood.

Agrobacteria are well-known as phytopathogenic bacteria responsible for crown gall disease affecting a wide range of plant species (Dessaux and Faure, 2018). These bacteria are only pathogenic when they carry a plasmid named Ti (for tumor inducing), the driver of agrobacteria pathogenesis (Hooykaas, 2023). Agrobacteria are also, and above all, ubiquitous soil borne and rhizospheric bacteria able to establish commensal or even beneficial interactions with plant roots (Costechareyre et al., 2010; Shams et al., 2012; Naqqash et al., 2016). Field investigations consistently showed that several species of agrobacteria generally co-exist in the same biotopes (bare soil, rhizosphere, host plants) (Nesme et al., 1987; Vogel et al., 2003; Shams et al., 2012; Bouri et al., 2016). This means that co-existing species must exploit different resources to avoid competitions with their closest relatives, and thus have at least partly different ecological niches. A better understanding of these ecological niches and interactions with plants could provide insights for combating this pathogen. In this sense, *Agrobacterium* species harbors species-specific genes (e.g. encoding specific ecological functions) potentially responsible for their adaptation to a species-specific plant ecological niche (Lassalle et al., 2011, 2017).

Comparative genomics performed by Lassalle et al. (2011) and strengthened by recent genomics analyses revealed that the species-specific genes of the genomic species *Agrobacterium fabrum* (*i.e. Agrobacterium* species genomovar G8) were mainly clustered into seven genomic islands on its genome, called "specific-regions" or SpG8 (for G8-specific coding DNA sequences), going from SpG8-1 to SpG8-7 (Du et al., 2023). Some specific-regions were subsequently divided into subclusters encoding homogeneous functions (**Table 2**). The annotation of these specific regions leans towards a close connection with the plant, since they encode coherent putative pathways related, for example, to the metabolism of trophic resources derived from plants (SpG8-1a, SpG8-1b, SpG8-4, SpG8-5), to the expression of environmental sensing systems (SpG8-7), or to the synthesis of extracellular compounds, such as siderophores (SpG8-3) or curdlan (SpG8-2a).

The latter, a glucose polymer induced under stress conditions, surrounds and protects bacteria against heat and desiccation (Kim et al., 1999; McIntosh et al., 2005; Matthysse, 2018). Even if the role of curdlan in the attachment of *Agrobacterium* to biotic or abiotic surfaces is not clear and not well documented, the genes involved in curdlan biosynthesis are upregulated when *A. fabrum* colonizes *Arabidopsis* (González-Mula et al., 2018) thus suggesting a correlation with biofilm implementation and development.

One well-studied specific-region, the SpG8-1b, encodes the complete degradation pathway of some hydroxycinnamic acid (HCA) that will ultimately be used as a source of carbon and energy by *A. fabrum* (Campillo et al., 2014; Meyer et al., 2018). HCA are common plant specialized metabolites released in soil during the decay of root cells and described as chemotactic signals for agrobacteria (Whitehead et al., 1983; Parke et al., 1987). This degradation pathway could lead to a competitive advantage of *A. fabrum* over other agrobacteria in HCA-enriched environments such as rhizosphere (Lassalle et al., 2011; Meyer et al., 2018).

Another well-known specific-region is the SpG8-3, that has been shown to encode the biosynthesis of siderophores and reuptake/regulatory system required for *A. fabrum* C58 growth under iron limiting conditions (Rondon et al., 2004). These siderophores, named fabrubactins A and B with unusual structures (Vinnik et al., 2021), could provide a fitness advantage to outperform competitors, especially in complex habitats like rhizosphere (Lassalle et al., 2011). A coordinated expression between SpG8-1b and SpG8-3 specific-regions has been observed, both induced in the presence of HCAs (Baude et al., 2016). The combined expression of these two specific regions suggests that several of them could be involved in the adaptation to a plant-related ecological niche (Lassalle et al., 2017).

Given this strong interconnection between *A. fabrum*-specific regions and the plant, they are thus suspected to have an effect on root during bacteria-plant interactions. The roots are highly colonized by a wide range of soil microorganisms, establishing different types of interactions which involve rhizosphere chemical dialogues via primary and specialized metabolites (Badri et al., 2009; Massalha et al., 2017). Indeed, plants can synthesize a wide diversity of specialized metabolites, which are known to be key components for the adaptation to abiotic and/or biotic conditions and notably plant microbiome (Bennett and Wallsgrove, 1994; Jan et al., 2021; Koprivova and Kopriva, 2022). Metabolomic studies have been carried out to identify metabolites involved in bacteria-plant interactions as it is a powerful approach to achieve an accurate comparative study of the metabolite content modulation (Gupta et al., 2022). Recent studies on the effect of phytobeneficial and phytopathogenic bacteria on plants (Miotto-Vilanova et al., 2019; Valette et al., 2019; Zeiss et al., 2019; Nong et al., 2023) highlighted significant changes on root metabolite profiles in particular phenolic compounds ranging from simple phenolic acids to complex polymerized tannins. These compounds are well described to play multifunctional roles in plant-microbial interactions notably signaling molecules for rhizobacteria including agrobacteria (Mandal et al., 2010).

To evaluate the involvement of *A. fabrum*-specific regions in the specific interaction between plant and this bacterial species, investigations of their impact on the plant and specifically on its specialized metabolites were performed. To this end, deletion mutant strains of each *A. fabrum*-specific region were inoculated on *Medicago truncatula* roots, which is frequently colonized by *Agrobacterium* and is a well-establishing plant model for studying plant-bacteria interactions (Cook, 1999; Farag et al., 2008). Root metabolite profiles were analyzed and compared by ultra–high-performance liquid chromatography coupled to a diode array detector and an electrospray ionization quadrupole time-of-flight mass spectrometer (UHPLC-UV/DAD-ESI-MS QToF). Highlighted discriminating metabolites were annotated or identified by tandem mass spectrometry (MS/MS). This led to a better understanding of the influence of specific bacterial genes on plant root metabolites. Almost all *A. fabrum-*specific regions were found to influence specialized root metabolites (mainly flavonoid classes), with different intensities, but in coordinated fashion, in response to the interaction of *A. fabrum* with *M. truncatula* roots. This work underlined the involvement of species-specific regions of *A. fabrum* in its interactions with plant roots, in particular via their influence on the phenolics of *M. truncatula* roots.

## MATERIALS AND METHODS

### Chemicals

Organic solvent for extraction was distilled methanol purchased from VWR Chemicals (Paris, France). Acetonitrile, water and acetic acid (Optima® UHLC-grade, Fisher Scientific, Geel, Belgium) were used for UHPLC-UV/DAD-ESI MS QTOF analyses. Tryptophan was obtained from Agilent Technologies (Waidbronn, Germany). Flavonoids (*i.e.* 7-4’-dihydroxyflavone, liquiritigenin, diosmetin, ononin, apigenin, kaempferol and kaempferol 3-O-glucoside) were obtained from Extrasynthese (Genay, France). Maesopsin 4-O-glycoside was kindly provided by Pr. Laurence Voutquenne-Nazabadioko from the Reims Institute of Molecular Chemistry (ICMR, UMR CNRS 7312).

### Bacterial strains, plasmids and culture conditions

The bacteria and plasmids used for this study are listed in **Table 1**. *Escherichia coli* was grown with shaking (160 rpm) at 37°C in LB medium. Growth media were supplemented with appropriate antibiotics (gentamicin 15 µg/ml, kanamycin 25 µg/ml) when necessary. The *A. fabrum* strains were grown at 28°C in YPG-rich medium (yeast extract, 5g.L^-1^; peptone, 10g.L^-1^; glucose, 10g.L^-1^; pH 7.2) liquid or solid when supplemented with agar (15g.L^-1^) or AB minimal medium supplemented with succinate (10mM) (Chilton et al., 1974). When necessary, growth media were supplemented with appropriate antibiotics (gentamicin 20 µg/ml, neomycin 25 µg/ml, kanamycin 25 µg/ml).

**Table 1.** Strains and plasmids used in this study.

| Strains | Relevant genotype and features | Reference |
| --- | --- | --- |
| ***Escherichia coli*** |  |  |
| JM109 | *endA1 glnV44 thi-1 relA1 gyrA96 recA1 mcrB+ Δ(lac-proAB) e14- [F'traD36 proAB+ lacIq lacZΔM15] hsdR17(rK-mK+)* | NEB catalog |
| ***Agrobacterium fabrum*** |  |  |
| C58 (WT1) | Wild-type | CFBP 1903 |
| C58 Kan (WT2) | pTiatu6148:Km, Neo/Kan<sup>R</sup> | (Lang et al., 2013) |
| C58ΔSpG8-1b (Δ1b) | C58 deleted of *atu1409 – atu1423* cluster; Neo/Kan<sup>R</sup> | (Lassalle et al., 2011) |
| C58ΔSpG8-2a (Δ2a) | C58 deleted of *atu3054 – atu3059* cluster; Neo/Kan<sup>R</sup> | (Lassalle et al., 2011) |
| C58ΔSpG8-2b (Δ2b) | C58 deleted of *atu3069 – atu3073* cluster; Neo/Kan<sup>R</sup> | Present work |
| C58ΔSpG8-3 (Δ3) | C58 deleted of *atu3663 – atu3693* cluster; Neo/Kan<sup>R</sup> | (Baude et al., 2016) |
| C58ΔSpG8-4 (Δ4) | C58 deleted of *atu3808 – atu3830* cluster; Neo/Kan<sup>R</sup> | Present work |
| C58ΔSpG8-5 (Δ5) | C58 deleted of *atu3947 – atu3952* cluster; Neo/Kan<sup>R</sup> | Present work |
| C58ΔSpG8-7a (Δ7a) | C58 deleted of *atu4285 – atu4294* cluster; Neo/Kan<sup>R</sup> | Present work |
| C58ΔSpG8-7b (Δ7b) | C58 deleted of *atu4295 – atu4307* cluster; Neo/Kan<sup>R</sup> | Present work |
| **Plasmids** |  |  |
| pGEM-T-Easy | Cloning vector for PCR fragments; Amp<sup>R</sup>; *lacZ* | Promega |
| pOT1e | Promoter-probe vector based on pBBR1MCS-5 replicon; contains promoterless eGFP and MCS between two transcriptional terminators; Gm<sup>R</sup> | (Allaway et al., 2001) |
| pJQ200sk | Suicide vector; P15A *sacB*; Gm<sup>R</sup> | (Quandt and Hynes, 1993) |
| pGEM-T *Patu1416* | Upstream region of *atu1416* gene cloned into pGEM-T-Easy; Amp<sup>R</sup> | Present work |
| pGEM-T *Patu3057* | Upstream region of *atu3057* gene cloned into pGEM-T-Easy; Amp<sup>R</sup> | Present work |
| pGEM-T *Patu3069* | Upstream region of *atu3069* gene cloned into pGEM-T-Easy; Amp<sup>R</sup> | Present work |
| pGEM-T *Patu3675* | Upstream region of *atu3675* gene cloned into pGEM-T-Easy; Amp<sup>R</sup> | Present work |
| pGEM-T *Patu3817* | Upstream region of *atu3817* gene cloned into pGEM-T-Easy; Amp<sup>R</sup> | Present work |
| pGEM-T *Patu3948* | Upstream region of *atu3948* gene cloned into pGEM-T-Easy; Amp<sup>R</sup> | Present work |
| pGEM-T *Patu4292* | Upstream region of *atu4292* gene cloned into pGEM-T-Easy; Amp<sup>R</sup> | Present work |
| pGEM-T *Patu4299* | Upstream region of *atu4299* gene cloned into pGEM-T-Easy; Amp<sup>R</sup> | Present work |
| pOT1e *Patu1416* | Upstream region of *atu1416* gene cloned into pOT1e; Gm<sup>R</sup> | Present work |
| pOT1e *Patu3057* | Upstream region of *atu3057* gene cloned into pOT1e; Gm<sup>R</sup> | Present work |
| pOT1e *Patu3069* | Upstream region of *atu3069* gene cloned into pOT1e; Gm<sup>R</sup> | Present work |
| pOT1e *Patu3675* | Upstream region of *atu3675* gene cloned into pOT1e; Gm<sup>R</sup> | Present work |
| pOT1e *Patu3817* | Upstream region of *atu3817* gene cloned into pOT1e; Gm<sup>R</sup> | Present work |
| pOT1e *Patu3948* | Upstream region of *atu3948* gene cloned into pOT1e; Gm<sup>R</sup> | Present work |
| pOT1e *Patu4292* | Upstream region of *atu4292* gene cloned into pOT1e; Gm<sup>R</sup> | Present work |
| pOT1e *Patu4298* | Upstream region of *atu4298* gene cloned into pOT1e; Gm<sup>R</sup> | Present work |

### Construction of the deletion mutants and transcriptional fusions

*A. fabrum* deletion mutants were constructed by recombination with the suicide vectore pJQ200sk vector according to a strategy previously described by Lassalle et al. (2011). Briefly, the recombinant region containing the flanking fragments of the sequence to be deleted (primers listed in **Table S1**) and a fragment encoding the *nptII* gene (neomycin/kanamycin resistance) amplified from plasmid KD4 (Datsenko and Wanner, 2000) were cloned into pJQ200sk (Quandt and Hynes, 1993) and then inserted into *A. fabrum* C58 by electroporation. Single-crossover integration was selected by kanamycin and neomycin resistance on YPG medium plates. Double crossover events were identified on YPG medium plates containing 10% sucrose. Deletion mutants were verified by diagnostic PCR with appropriate primers (**Table S1**) and DNA sequencing (GenoScreen, Lille, France).

The eGFP transcriptional fusions were obtained as follows. The promoter regions of the targeted genes in each *A. fabrum* specific region were PCR amplified with specific primers listed in **Table S1**. Then, PCR fragments obtained were cloned into the pGEM-T Easy vector (Promega, Madison, WI) according to manufacturer’s instructions. After digestion of the resulting plasmids with HindIII and SalI, fragments were subcloned into pOT1e (Allaway et al., 2001) digested with the same enzymes. Finally, transcriptional reporter constructions were inserted into C58 by electroporation, gentamicin resistant colonies were selected. Constructions were verified by diagnostic PCR using pOT1eF and pOT1eR primers listed in **Table S1** (Pothier et al., 2007).

### Phenotypic characterization

C58 wild-type and mutants for the specific-regions were subjected to phenotypic characterization using Biolog phenotype MicroArrays PM1 and PM2A (Biolog Inc, Hayward, CA, United States). Briefly, all strains were grown on YPG medium for 48h at 28°C and bacteria were collected from the agar surface and suspended in inoculating fluid (IF-0) to a cell density of 42% transmittance on a turbidimeter (Biolog, Inc., Hayward, CA, USA). 5 ml of cell suspension were added to 25 ml of IF-0+dye MixA to obtain a final transmittance of 85%. Each well of PM1 and PM2A plates were inoculated with 100 µl of this suspension. Plates were incubated at 28°C in the Omnilog for 60 hours with readings taken every 20 minutes. After incubation, the Omnilog ™ system was used to measure carbon substrate utilization. Biolog phenotypic MicroArray analyses were performed independently twice.

### Plant material and *in vitro* culture procedures

All plant experiments were conducted on *Medicago truncatula* Gaertn (cultivar Jemalong line A17). *M. truncatula* seeds were surface disinfected by soaking in a 3.5% sodium hypochlorite solution before being thoroughly rinsed five times with sterile distilled water. To vernalize the seeds, they were put in the dark at 4°C overnight before use. The seeds were then scarified and placed in square Petri dishes (120 mm x 120 mm) containing 0.8% agar plant medium (A7921, Sigma Aldrich, St Louis, MO, USA) supplemented with nutrient solution (15% nitrogen 15% potassium, 30% phosphorus at 1.5 g/L, Plant-Prod 15-15-30, Fertil s.a.s., Le Syndicat, France). All Petri dishes were placed vertically, and the lower part was covered to protect the root development from light. The plants were incubated for 14 days in a phytotron with 16 hours light daily and a temperature between 24°C and 28°C.

### Confocal microscopy observation

For confocal observation, bacterial cells harboring the eGFP transcriptional fusions constructed were inoculated on the *M. truncatula* seeds with 10 µL of overnight culture (2x10^5^ bacteria.mL^-1^). At 14 dpi, *M. truncatula* roots were mounted between a slide and a coverslip in a commercial mounting fluid (Aqua Poly/Mount, Polysciences, Inc., Warrington, PA). Microscope observation was performed using a confocal laser scanning microscope (LSM 800 Meta Confocal Microscope, Zeiss, Oberkochen, Germany). eGFP was excited with an argon laser at 488 nm, and fluorescence was monitored at 528 nm. Analyses of images (five plants per condition) were performed with the LSM 800 software (Zeiss, Oberkochen, Germany).

### Bacterial root colonization

Before inoculation with *A. fabrum*, scarified seeds were placed on plates containing 0.8% agar plant cell culture for 2 days in the darkness. Seedling were inoculated with 5 µL of bacterial culture (prepared from overnight culture in YPG, washed with NaCl 0.8 % and resuspended at 10^5^ cfu/ml in NaCl 0.8%) from *A. fabrum* wild-type and mutants. For each strain, concentration of the inoculum was verified by CFU determination on YPG agar medium. Petri dishes with plants were placed vertically and the lower part was covered to protect the root development from light. The plants were incubated for 15 days in a phytotron with 16 hours light daily and day/night temperature 20°C/ 18°C. To determine bacterial colonization level, roots were ground at 14 dpi. Serial dilutions were plated on YPG medium, colonies were counted after 2 days of incubation at 28°C. Bacteria abundance was expressed as log10 CFU per gram of root dry weight.

### Metabolomic study of plant-bacteria interaction Plant bacteria co-culture procedures

To perform the metabolomic study, *Agrobacterium* strains were inoculated in the supercooled agar at a concentration of 1x10^7^ bacteria.mL^-1^ of agar plant medium before seedling. One group of *M. truncatula* seeds was cultivated without bacterial inoculation to carry out the non-inoculated (NI) condition as a control. All petri dishes were placed vertically, and the lower part was covered to protect the root development from light. The plants were incubated for 14 days in a phytotron with 16 hours light daily and a temperature between 24°C and 28°C.

### Roots harvesting and extraction of phenolic compounds

After 14 days of incubation, the roots were harvested, pooled, frozen in liquid nitrogen (metabolism quenching) and freeze-dried. They were then grinded using a bead mill TissueLyser II system (Qiagen, Hilden, Germany). Four or five replicates were performed per condition, each replicate consisting of root pools of several seedlings. Dried powders were first extracted twice with methanol 80% and then extracted twice with pure methanol with homogenization and 10 min of sonication at room temperature. The combined extracts were then centrifuged (10 min, 19 745 × g) and the supernatants, particle free, were evaporated to dryness at 30°C in a Centrivap concentrator (Labconco, USA) to constitute the dried crude extracts. These extracts were weighed, dissolved in methanol 60% at 10 mg/mL and conserved at -80°C prior to analysis. A quality control (QC) sample was prepared containing 20µL of each extract. A bacterial extract was also prepared following a similar extraction process applied to bacterial culture performed without plant, which constitutes another type of control.

### Metabolites analysis by UHPLC-UV/DAD-ESI-MS QTOF

Metabolites analysis was performed on an UHPLC Agilent 1290 coupled to a UV-vis Diode Array Detector (Agilent 1290 Infinity series) and an Accurate-Mass Q-ToF 6530 spectrometer (Agilent Technologies). Liquid chromatography was carried out using a Nucleodur Sphinx RP C_18_ column (2mm × 100mm, 1.8µm, Macherey-Nagel) maintained at 50°C with an injection volume of 3 µL of sample.

The mobile phase was a mixture of acetonitrile and acidified UHPLC-grade water (0.4% acetic acid) applied at a flow-rate of 0.5mL/min, and with the following gradient (acidified H_2_O:CH_3_CN, [v/v]) starting at 98:2 for 1.5min, increasing to 76:24 at 24.5min, going to 0:100 in 4min and maintained for 2min, before returning to the starting conditions in 1min and equilibrating for 2min. The QTOF-MS instrument was operated for MS analyses under the following conditions: the ion source ESI (Agilent Jet Stream thermal gradient focusing Technology) in positive ionization mode, with a 320°C gas temperature, a 11L/min gas flow, a nebulizer pressure at 40 psi, a 360°C sheath gas temperature with 12L/min flow rate, and with the capillary, nozzle, and fragmentor voltages at 3,000 V, 500 V, and 150 V, respectively. The acquisition mass range was from *m/z* 100 to 2000. The QC sample was initially analyzed and then injected after every 8-9 samples in the run sequence to monitor the repeatability of the analysis. For characterization of discriminating compounds, complementary MS and tandem MS experiment were performed in positive and negative ionization modes and with different collision energies (10, 20, 30 or 40V) in AutoMS/MS mode. The UHPLC-UV/DAD-ESI-MS QTOF device was managed by the Mass Hunter Workstation Acquisition B.07.00 software and the resulting data was reprocessed with the Mass Hunter Qualitative Analysis B.07.00 software (Agilent Technologies). Identification or annotations of discriminating metabolites were performed by the analysis of the UV-vis spectrum and the positive and negative MS and MS/MS spectra. These data were compared to in-house database (SeMetI), and to literature data, first on compounds already described in *M. truncatula* but also on chemical data of metabolites from other plants. When possible, comparisons with standards (the entire molecule or the aglycone moiety) allowed confirming metabolite identity or at least the aglycone nature.

### Metabolite profiling data treatment and statistical analyses

First, integration of each peak of the UV-chromatograms at 280nm (wavelength allowing the detection of a wide range of secondary metabolites and especially phenolic compounds) was performed. The retention time alignment between all samples was made for each peak, resulting in a metabolite matrix with samples in rows and absolute areas (A_abs_) of the 92 detected metabolites in columns. Then for each metabolite within a given sample, its relative peak area (*i.e.* relative abundance in the sample= A_rel_) was calculated over the total area of the sample chromatogram (ΣA_abs_) as follows: %A_rel_ = A_abs_ / ΣA_abs_ × 100. In the resulted metabolite matrix, only the most abundant metabolites (relative abundance >1%) were considered to continue the study (28 peaks). Partial least squares discriminating analysis (PLS-DA) was made from this matrix to visualize the global effect of bacterial inoculation (WT strain, mutant strains and the NI condition) on root phenolic profiles of *M. truncatula*. Principal Component Analyzes (PCAs) were made between the WT strain condition and each mutant strain condition to stand out more precisely the effect of each *A. fabrum* specific-regions on root secondary metabolite profiles. The statistical study was oriented to a t-test to compare each condition with the WT strain condition to highlight significantly different metabolites. Statistical analyses were performed using R software (R Studio version 0.99.491).

## RESULTS

To evaluate the influence of specific regions on root metabolites during the *A. fabrum-M. truncatula* interaction, mutant strains with precise deletion in the specific-regions (*i.e.* specific genes of *A. fabrum* species) were previously obtained or in this study (**Table 1 and 2**). All the genetic constructions were performed in the model strain *A. fabrum* C58.

**Table 2.**
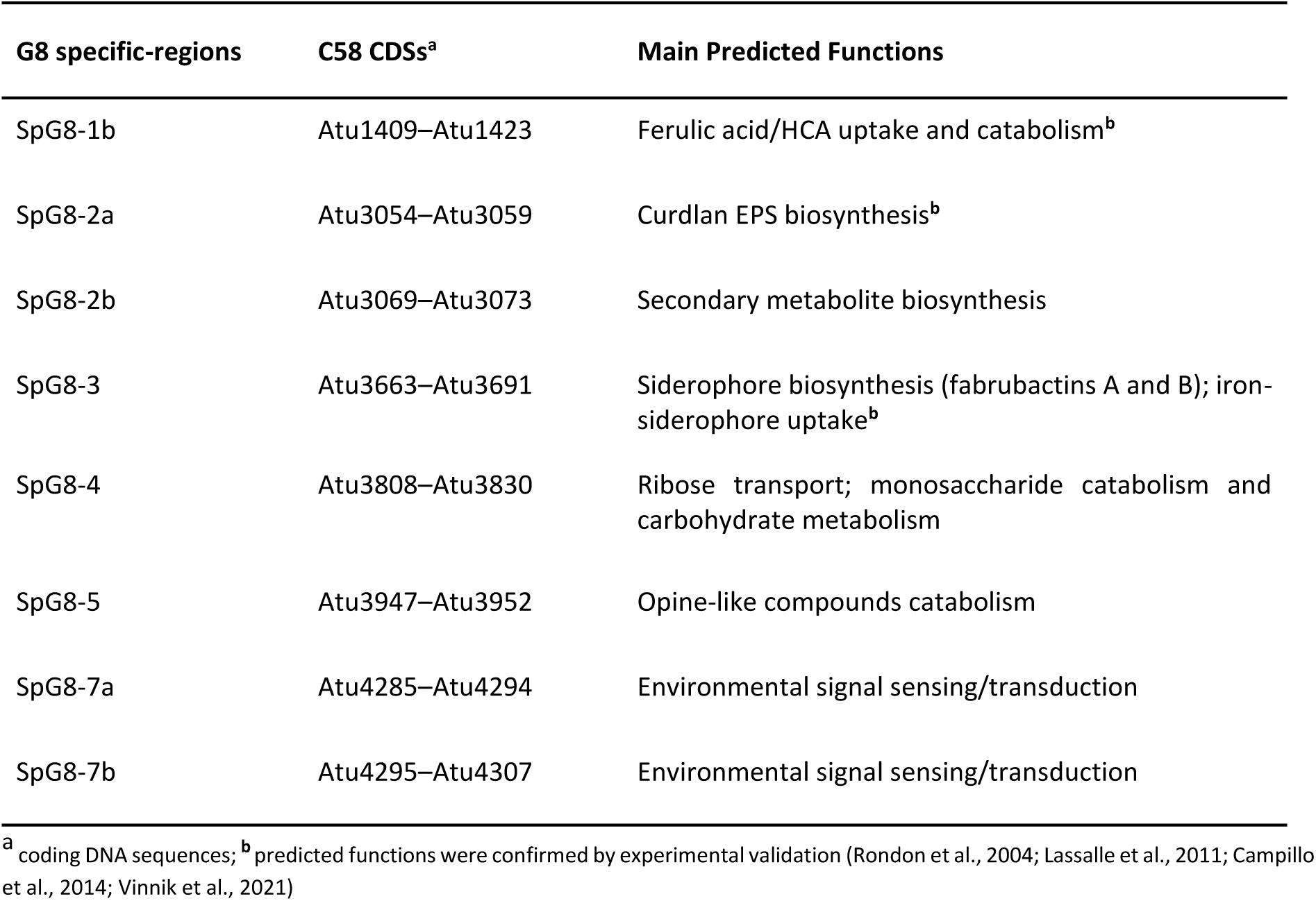
Characteristics and function annotations of *A. fabrum* gene specific regions (Lassalle et al., 2011)

### 1. Phenotypic characterization and root colonization capacity of mutants in the specific-regions

First, no significant difference in growth rate was observed *in vitro* (in rich medium and minimal medium) between wild-type C58 and the deletion mutant strains, or between mutant strains (data not shown). Biolog phenotype MicroArrays were used to further explore the functions of specific-regions. All but three deletion mutant strains showed the same use of carbon sources than the wild-type strain. C58ΔSpG8-3, C58ΔSpG8-5, and C58ΔSpG8-7a mutant strains are different from the wild-type strain in their use of several hexuronic acid substrates (**Fig. S1**). For instance, a decrease of metabolic activity of the three mutant strains was observed in the presence of D-galacturonic and D-gluconic acids, the most important decrease was observed in the presence of gluconic acid for C58ΔSpG8-3 and of galacturonic and glucuronic acids for C58ΔSpG8-5.

In a second step, the non-detrimental effect of these mutations on the root colonization was verified. Wild-type C58 and all the deletion mutant strains of *A. fabrum* were singly inoculated on *M. truncatula* germinating seeds and their abundance on roots was determined at 15 dpi. Some similar levels of colonization (mean values from 9.0 to 10.3 log10 CFU per gram of root dry weight) were observed between the wild-type and the deletion mutant strains (**Fig. S2**). Thus, these *A. fabrum*-specific region deletions have not been detrimental for the establishment and the survival of the mutant bacterial strains on the roots of *M. truncatula* seedlings *in vitro*-cultivated.

### 2. All *A. fabrum* specific regions are expressed on *M. truncatula* roots

The expression of all these specific-regions on *M. truncatula* roots were also validated with eGFP transcriptional fusion. For each specific-region, one promoter was selected based on annotation of genes and used for transcriptional fusion construction with eGFP as reporter gene (**Table 1**). Each transcriptional fusion was introduced in the wild-type C58 strain and inoculated on *M. truncatula* seedlings. Root microscopic observations on five plants per condition at 14 dpi showed a green fluorescence of each reporter bacteria (**Fig. S3**). All transcriptional fusions in the WT strains are induced in contact with the *M. truncatula* root system. This result reveals that for all the *A. fabrum*-specific regions at least one operon was activated during its interaction with *M. truncatula* roots.

### 3. Effect of *A. fabrum* wild-type or mutant strains inoculation on *M. truncatula* seedlings

All these verified *A. fabrum* strains (wild-type C58 and deletion mutant strains) were individually inoculated onto the root of *M. truncatula* germinating seeds to evaluate their impact on the plant compared to uninoculated plants. After 14 days of *in vitro* co-culture, *M. truncatula* roots and aerial parts were separately harvested and dried. Whether *M. truncatula* seedlings were non-inoculated or inoculated with the wild-type C58 strain or the deletion mutant strains, similar dry matters were obtained from roots (mean values ± SE, from 0.65 ± 0.04 to 1.00 ± 0.15 mg dry root per plant) and from aerial parts (mean values ± SE, from 2.43 ± 0.07 to 3.29 ± 0.33 mg dry aerial part per plant, **Fig. 1**). At 14 dpi under our conditions, inoculation of seedlings with *A. fabrum* strains (wild-type C58 or deletion mutants) did not affect *M. truncatula* development in terms of root and aerial dry matter.

**Fig 1.**
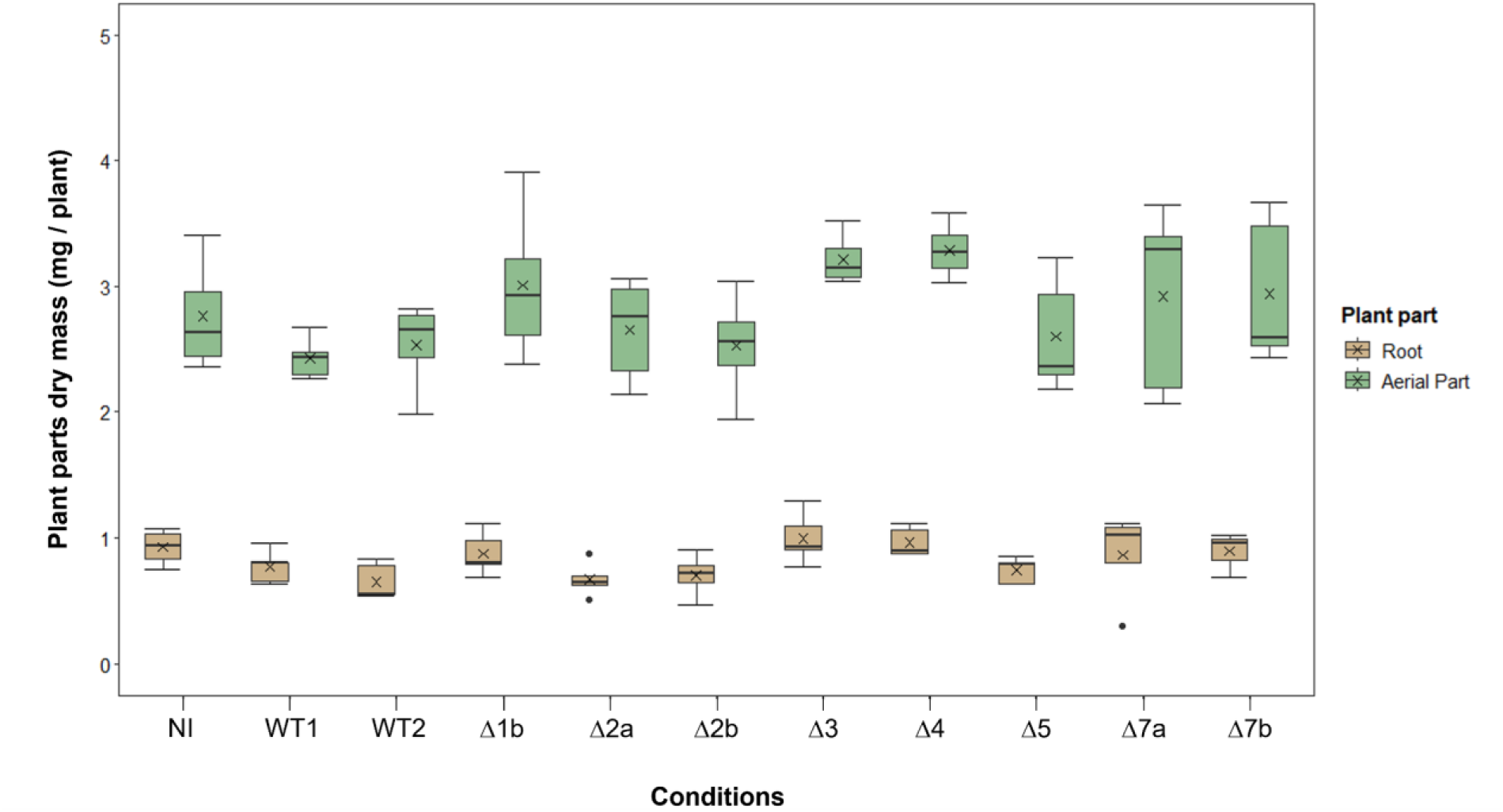
Boxplots illustrating the dry mass of roots and aerial parts from *Medicago truncatula* inoculated or not by *A. fabrum* wild-type or deletion mutant strains. For each condition at 14 dpi, the roots and aerial parts of *M. truncatula in vitro* cultivated were harvested and dried. Dry masses of these plant parts were weighed and expressed as mg per seedling. Boxes cover 50% of the data. Central lines represent the medians and whiskers represent the minimum and maximum values among non-atypical data. The cross (×) denotes the mean value of the data (n=4 or 5). Similar dry masses were obtained on the one hand for roots and on the other for aerial parts, whether *M. truncatula* seedlings were non-inoculated (NI) or inoculated with the wild-type (WT1, WT2) or the deletion mutant strains (Δ1b = C58ΔSpG8-1b; Δ2a = C58ΔSpG8-2a; Δ2b = C58ΔSpG8-2b; Δ3 = C58ΔSpG8-3, Δ4 = C58ΔSpG8-4; Δ5= C58ΔSpG8-5; Δ7a = C58ΔSpG8-7a; Δ7b= C58ΔSpG8-7b). No statistically significant difference was observed for each plant part (Dunn’s test pairwise comparisons using p-values adjusted with the Bonferroni method, for roots of all conditions *p* > 0.322, for aerial parts of all conditions *p* > 0.285).

### 4. Influence of *A. fabrum* inoculation on root specialized metabolites of *M. truncatula*

At 14 dpi, the harvested and dried roots of *M. truncatula* were extracted with methanol (80% and 100%). Whether *M. truncatula* seedlings were non-inoculated or inoculated with the wild-type C58 strain or the deletion mutant strains, similar amounts of crude extract were obtained from roots (mean values ± SE, from 0.21 ± 0.01 to 0.33 ± 0.08 mg of crude extract per mg of root dry mass, **Fig. S4**).

The specialized metabolite profiles of *M. truncatula* roots between plants inoculated with wild-type C58 strain (WT condition) and non-inoculated plants (NI condition) or plants inoculated with deletion mutant strains of *A. fabrum* (mutant strain conditions) were compared. Root methanolic extracts were analyzed to obtain chromatograms at 280 nm, a wavelength allowing the detection of a wide range of plant specialized metabolites and especially phenolics compounds. The quality control (QC) analysis led to the detection of 92 peaks (**Fig. S5**), among which those with relative abundances higher than 1% were considered in this study. The variations in chromatographic profiles observed between conditions lie in the relative abundance of peaks rather than in the appearance or disappearance of peaks. Chromatographic profiles were visualized globally using Partial Least Squares Discriminant Analysis (PLS-DA) (**Fig. 2**) and precisely compared using Principal Component Analysis (PCA) (**Fig. 3**).

**Fig. 2.**
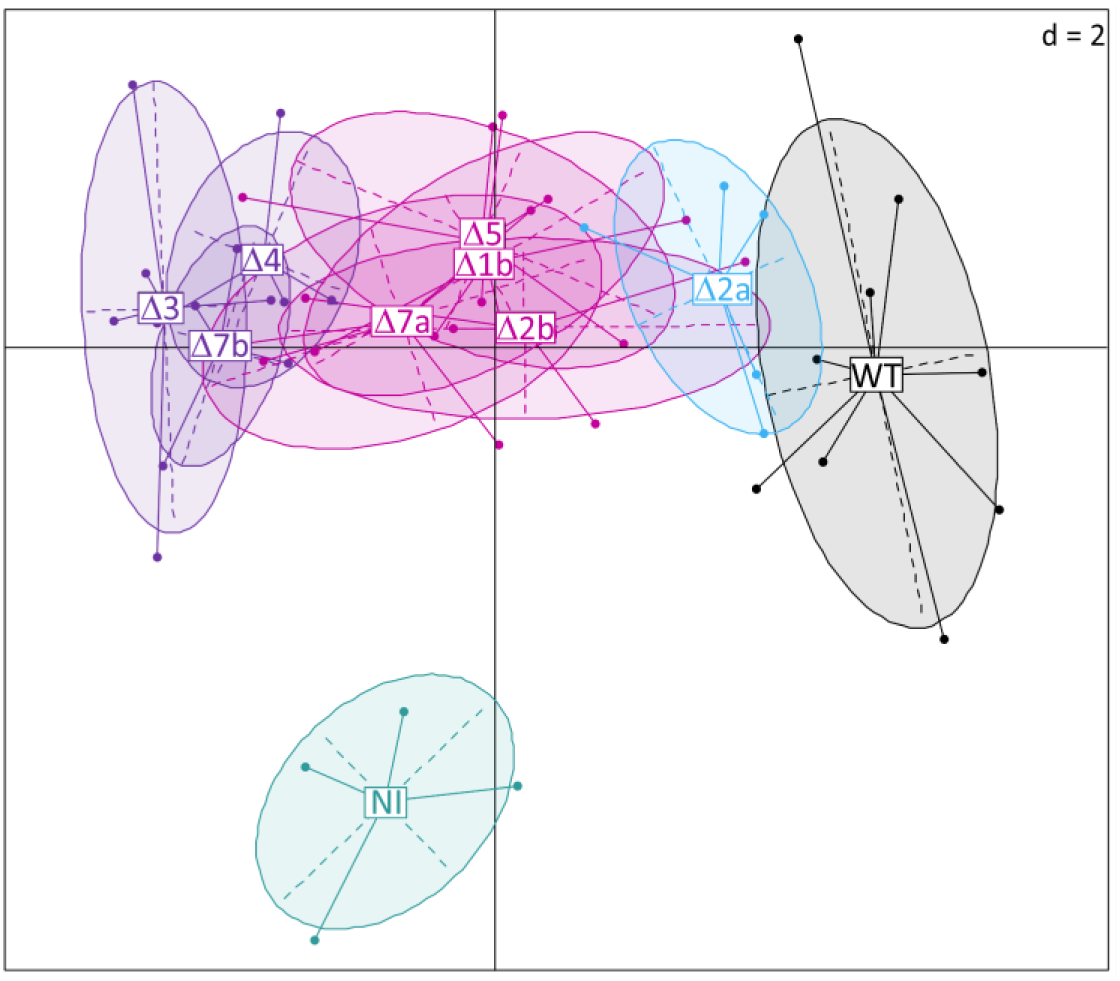
PLS-DA score plot of root specialized metabolites profiles of *M. truncatula* seedlings non-inoculated (NI) or inoculated with wild-type (WT) or mutant (Δx) strains of *A. fabrum* C58. PLS-DA were performed on the data matrix from chromatographic data (280 nm) obtained for each methanolic extract of *M. truncatula* roots based on peak areas and retention time (peaks > 1%). Plants were inoculated or not with *A. fabrum* C58 wild-type or deletion mutant strains of *A. fabrum*-specific regions. WT: plants inoculated with the wild-type strain (in black), NI: plants non-inoculated (in cyan), Δx: plants inoculated singly with each of the deletion mutant of *A. fabrum-*specific regions (for example Δ2a corresponds to plants inoculated with the C58ΔSpG8-2a mutant strain). In blue are drawn the closest root metabolite profile to the one of WT condition, in purple the further ones, and in pink the intermediate profiles. Δ1b = C58ΔSpG8-1b; Δ2a = C58ΔSpG8-2a; Δ2b = C58ΔSpG8-2b; Δ3 = C58ΔSpG8-3; Δ4 = C58ΔSpG8-4; Δ5= C58ΔSpG8-5; Δ7a = C58ΔSpG8-7a; Δ7b= C58ΔSpG8-7b.

**Fig. 3.**
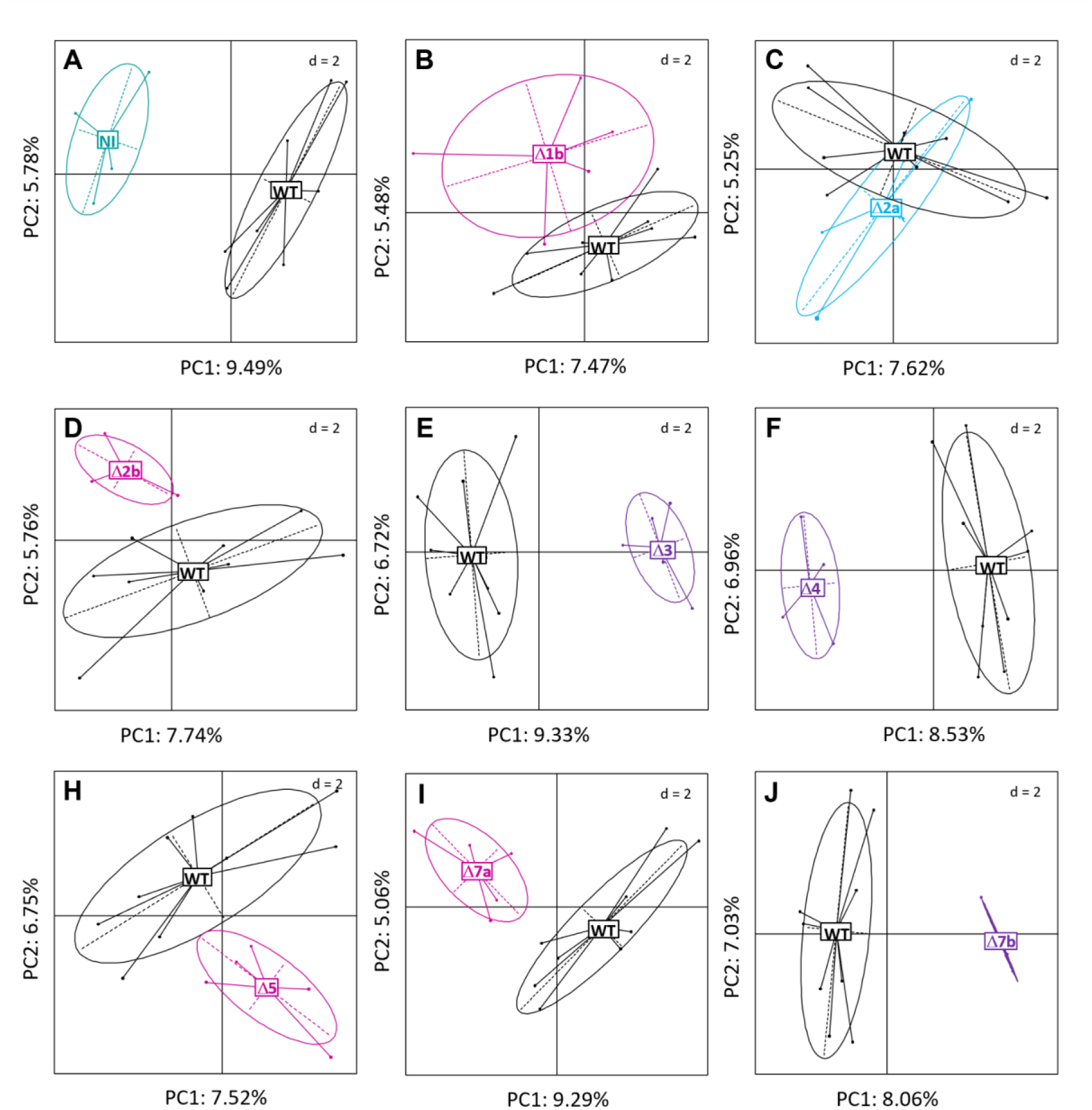
PCA score plots representing the comparison of root specialized metabolites profiles of *M. truncatula* seedlings between plant inoculated with wild-type strain (WT) and plant non-inoculated (NI) or inoculated with mutant (Δx) strains of *A. fabrum* C58. PCAs were performed on chromatographic data at 280 nm obtained for each methanolic extract of *M. truncatula* roots based on peak areas and retention time (peaks > 1%). Plants were inoculated or not with *A. fabrum* C58 wild-type or deletion mutant strains of *A. fabrum*-specific regions. WT: plants inoculated with the wild-type strain (in black), NI: plants non-inoculated (in cyan), Δx: plants inoculated singly with each of the deletion mutant of *A. fabrum-*specific regions (for example Δ2a corresponds to plants inoculated with the C58ΔSpG8-2a mutant strain). In blue are drawn the closest metabolites profiles to the one of WT, in purple the further ones, and in pink the intermediate profiles (color similar to that used in Fig. 2). Δ1b = C58ΔSpG8-1b; Δ2a = C58ΔSpG8-2a; Δ2b = C58ΔSpG8-2b; Δ3 = C58ΔSpG8-3; Δ4 = C58ΔSpG8-4; Δ5= C58ΔSpG8-5; Δ7a = C58ΔSpG8-7a; Δ7b= C58ΔSpG8-7b.

Two conditions corresponding to *A. fabrum* WT strain were used in this study (named WT1 and WT2): one was a native *A. fabrum* wild-type strain (WT1) and the second (WT2) had the *ntpII* kanamycin resistance gene used to construct deletion mutants. A first comparison by PCA did not show any separation of *M. truncatula* root metabolite profiles between these two WT conditions (**Fig. S6**). The statistical analyses on each peak highlighted only one (peak **39**) significantly different in its relative abundance between WT1 and WT2. As this peak was influenced by the presence of the *ntpII* gene, it was excluded from the data matrix for further analysis. The two wild-type conditions were then combined into a single condition, henceforth referred to as the WT condition.

#### 4.1. Global visualization of the root specialized metabolites profiles

The PLS-DA performed on all root metabolite profiles provided a global visualization (**Fig. 2**). On the one hand, the root metabolite profile of the NI condition seemed to be distinct from all the other profiles of the inoculated conditions. On the other hand, a gradient of root metabolite profiles seemed to appear between plants inoculated by WT C58 strain and deletion mutants of *A. fabrum*-specific regions strains (**Fig. 2**). The root metabolite profile of plants inoculated with mutant strain C58ΔSpG8-2a seemed to be the closest to that of the WT condition, while that of plants inoculated with mutant strain C58ΔSpG8-3 to be furthest apart. However, as it describes a large number of conditions for a relatively small number of variables, this PLS-DA representation cannot be considered as a predictive model.

#### 4.2. Evidence of bacterial effects on root specialized metabolites content

The PCA comparison of root metabolite profiles of the WT and NI condition showed discrimination between the two along PC1 (**Fig. 3A**). Eleven root metabolites significantly varied in their relative abundances between both conditions. Five of which are underabundant (**8**, **40**, **44**, **87** and **90**), the remaining six are overabundant (**24**, **25**, **26**, **36**, **65** and **79**) in the NI condition compared to the WT condition (**Fig. 4**). Among them, two compounds (**8** and **25**) were exclusively discriminating between the NI and the WT condition (*i.e.* not discriminating for the mutant strain conditions).

**Fig. 4.**
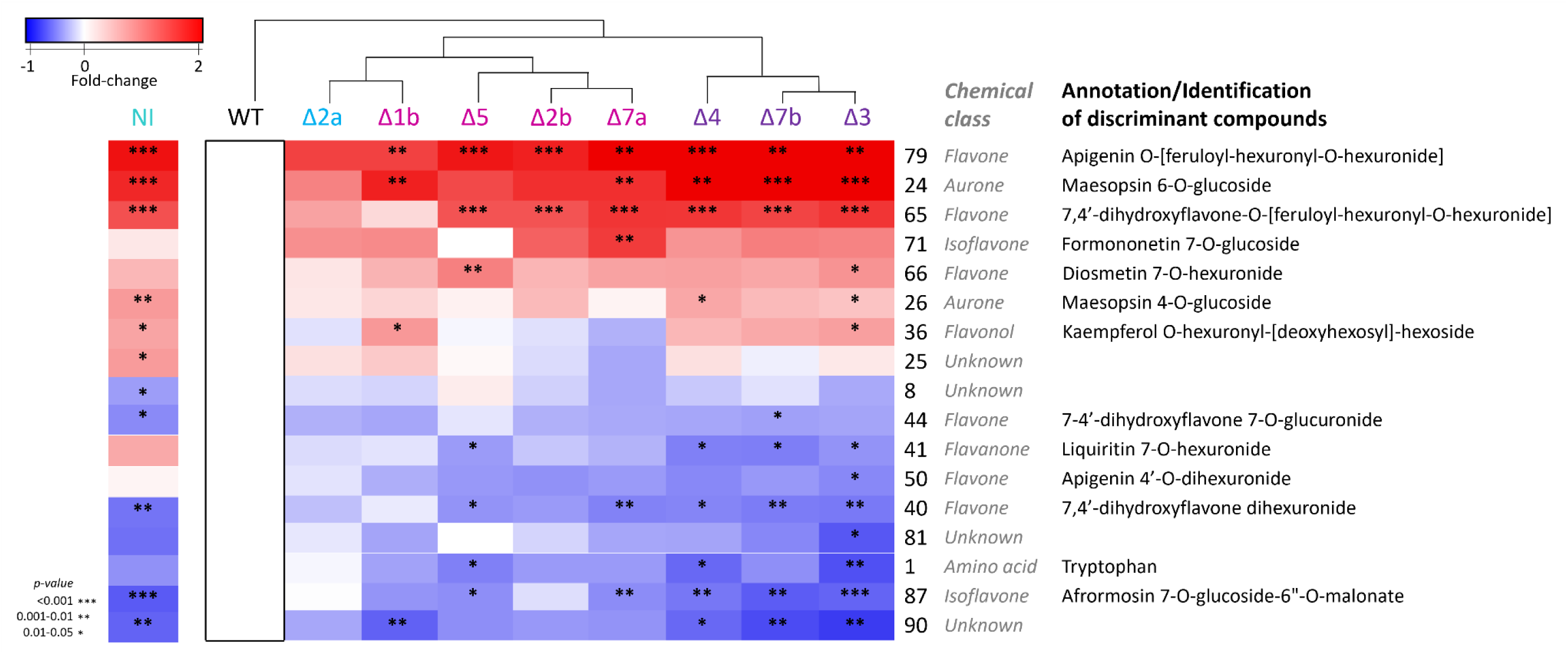
Heatmap of significantly discriminant metabolites according to their relative abundance in the roots of *Medicago truncatula* between plant inoculated with wild-type strain (WT) and plant non-inoculated (NI) or inoculated with mutant (Δx) strains of *A. fabrum* C58. Metabolites are represented as numbers on the right-hand side of the plot, using the metabolite number obtained from the data processing. Chemical class and annotation/identification are indicated when determined. These compounds are over-abundant (red) or under-abundant (blue) in plants either not inoculated (NI) or inoculated with each of the deletion mutant strains of *A. fabrum*-specific regions (Δx) compared to plants inoculated with C58 wild-type (WT) strain of *A. fabrum* used as reference. The colors of the mutant strains are similar to those used in Fig. 2 **and** Fig. 3. The *A. fabrum*-specific regions deleted in the mutant strains Δx and their predicted functions are as follows : Δ1b = C58ΔSpG8-1b (Ferulic acid/HCA uptake and catabolism); Δ2a = C58ΔSpG8-2a (curdlan EPS biosynthesis); Δ2b = C58ΔSpG8-2b (secondary metabolite biosynthesis); Δ3 = C58ΔSpG8-3 (siderophore biosynthesis - fabrubactins A and B; iron-siderophore uptake), Δ4 = C58ΔSpG8-4 (Ribose transport; monosaccharide catabolism and carbohydrate metabolism); Δ5= C58ΔSpG8-5 (opine-like compounds catabolism); Δ7a = C58ΔSpG8-7a (environmental signal sensing/transduction); Δ7b= C58ΔSpG8-7b (environmental signal sensing/transduction). Data were compared by analysis of variances between WT condition and the others one by one. The statistically significant differences are indicated with the symbol * (*** for *p-value* < 0.001; ** for 0.001< *p-value* <0.01; * for 0.01< *p-value* <0.05)

#### 4.3. Effect of A. fabrum specific regions on root specialized metabolite profiles

Root metabolite profiles of each mutant strain condition were singly compared with the WT condition (**Fig. 3B-J**). All root metabolite profiles from mutant strain conditions clearly separated from that of the WT condition, except for the C58ΔSpG8-2a mutant strain (**Fig. 3C**). Fifteen chromatographic peaks were found to display a significantly different relative abundance compared to the WT condition for at least one mutant condition (**Fig. 4**). Remarkably, there were only two patterns of change in metabolite relative abundance: on the one hand, compounds that were exclusively overabundant in mutant strain conditions compared to the WT condition (**24**, **26**, **36**, **65**, **66**, **71** and **79**), and on the other, compounds that were exclusively underabundant in mutant strains conditions (**1**, **40**, **41**, **44**, **50**, **81**, **87** and **90**) (**Fig. 4**). These analyses confirmed a gradient of differences in root metabolite profiles depending on the mutant strain conditions compared to the WT condition (**Fig.3**, **Fig. 4**). This gradient ranged from a non-different profile for plants inoculated with strain C58ΔSpG8-2a (**Fig. 3C**, **Fig. 4**), to profiles clearly differentiated from the WT condition (for plants inoculated with strains C58ΔSpG8-3, C58ΔSpG8-4 and C58ΔSpG8-7b) by variations in the relative abundance of 8 to 13 metabolites (**Fig. 3E-F-J, Fig. 4**). The intermediate root metabolite profiles (for plants inoculated with strains C58ΔSpG8-1b, C58ΔSpG8-5, C58ΔSpG8-2b and C58ΔSpG8-7a) showing significant differences in relative abundance of only 2 to 7 metabolites (**Fig. 3B-D-H-I, Fig. 4**).

### 5. Identification or annotation of discriminating metabolites

The UHPLC-UV/DAD-MS/MS QTOF data were explored to identify the discriminating compounds highlighted by statistical analyses. Study of the spectral data (UV-vis maxima; accurate mass; MS and MS/MS in positive and negative ionization mode) allowed the annotation and/or the identification of 13 compounds by comparison to bibliographical data and analyses of standard compounds when available (the entire molecule or the aglycone moiety). Chemical data of the annotated discriminating compounds are shown in **Table 3**. The detailed interpretation of these spectral data for the structural elucidation of these compounds were given in the supplementary materials (**supM1**). All but one discriminating metabolite are phenolic compounds belonging to the flavonoid family of different subclasses (aurone, flavanone, isoflavone, flavone and flavonol). The other is an amino acid (**1**, tryptophan), whose underabundance was significant in some conditions (**Fig. 4**). Based on their accurate mass, all the discriminating plant metabolites were searched in the control bacterial extracts. None of these metabolites was detected (data not shown). Likewise, the bacterial metabolites found in the bacterial extract were not detected in the obtained root plant extracts.

**Table 3.**
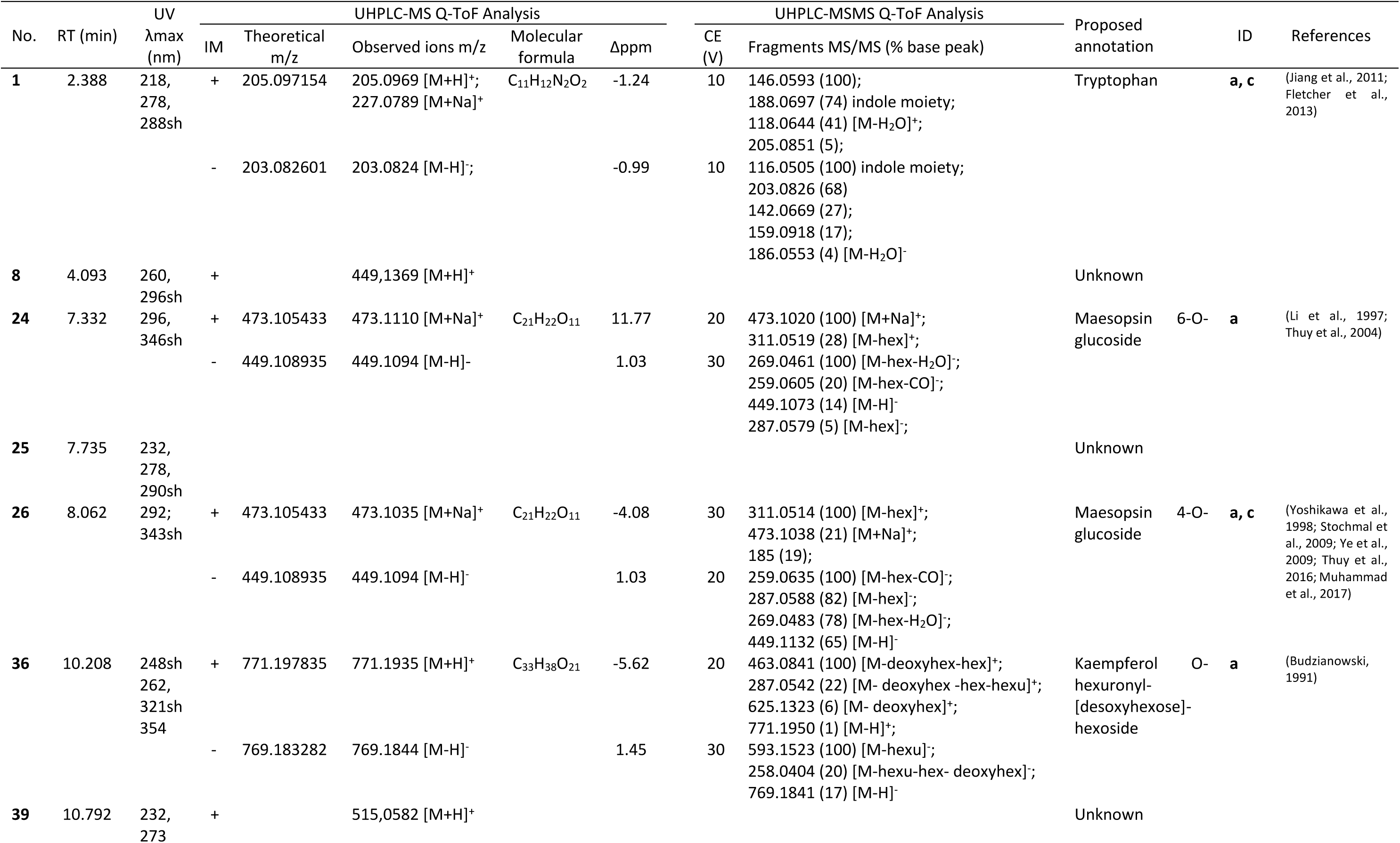
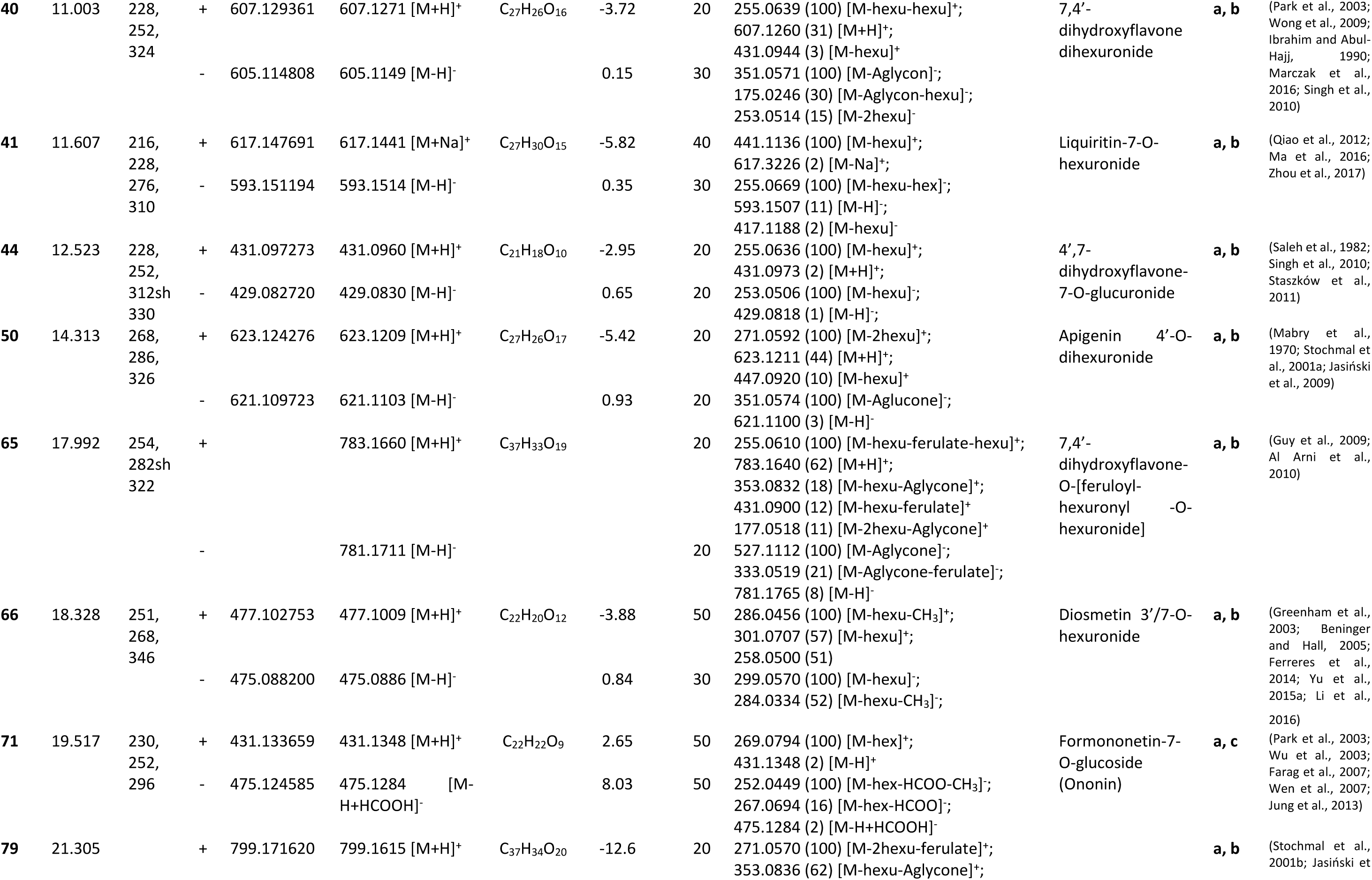
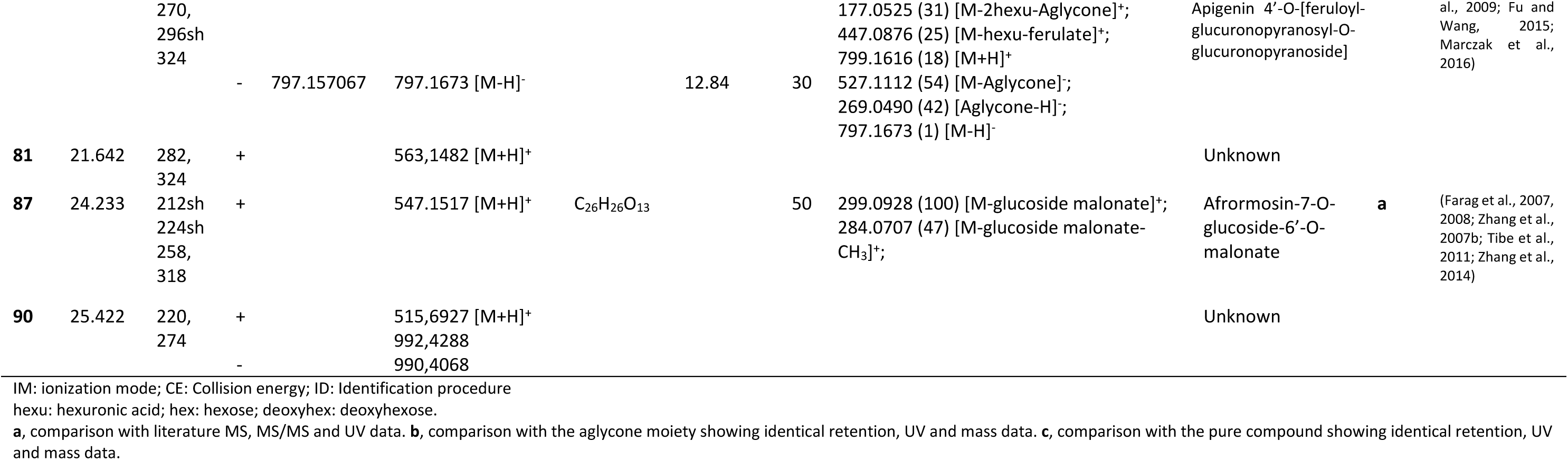
UPHLC-UV/DAD-ESI-MSMS Q-TOF data of discriminant metabolites in *M. truncatula* root in interaction with *A. fabrum* strains.

## DISCUSSION

Phenolic compounds are often involved in plant-bacteria interactions (Wang et al., 2022; Singh et al., 2023; Wang et al., 2023) and their content has been already shown to be affected in plant response to phytobeneficial and phytopathogenic bacteria (Walker et al., 2011; Miotto-Vilanova et al., 2019; Valette et al., 2019; Zeiss et al., 2019; Nong et al., 2023). In this context, this study focused on phenolic compounds using UHPLC-UV/DAD-ESI-MS QTOF analyzes to identify plant compounds whose relative abundances were significantly altered in roots inoculated with wild-type or mutant bacterial strains for the *A. fabrum*-specific gene regions. The aim was to assess the involvement of *A. fabrum*-specific regions in the interaction between the plant and this bacterial species, and more particularly their influence on plant specialized metabolites.

The annotation of the functions of these specific gene regions (**Table 2**) suggested a close connection with the plant (Lassalle et al., 2011). Here, eGFP transcriptional fusion experiments demonstrated that for all these *A. fabrum*-specific gene regions at least one operon was activated during its interaction with *M. truncatula* roots (**Fig.S3**), further supporting this hypothesis of a role for these genes in the bacterium’s interaction with the plant. To go further in the exploration of these specific gene regions, the putative interactions among their encoded proteins were investigated by applying the STRING v12 database. The STRING database (https://string-db.org/) systematically collects and integrates protein-protein interactions-both physical interactions as well as functional associations (Szklarczyk et al., 2023). The protein–protein interaction network of the *A. fabrum* C58 species specific proteins is presented in **Fig. 5**. All but SpG8-2a species specific regions had proteins linked to at least one protein from another species-specific region, suggesting a possible biological connection between them. The SpG8-3 protein cluster appears to be linked to the ones of four different species-specific regions (SpG8-1b, SpG8-2b, SpG8-5 and SpG8-7b). The connection between the specific regions SpG8-3 and SpG8-1b has already been experimentally validated. Indeed, coordinated expression has been observed after induction with hydroxycinnamic acids (HCAs) and the presence of one is required for expression of the other and vice versa (Baude et al., 2016).

**Fig. 5:**
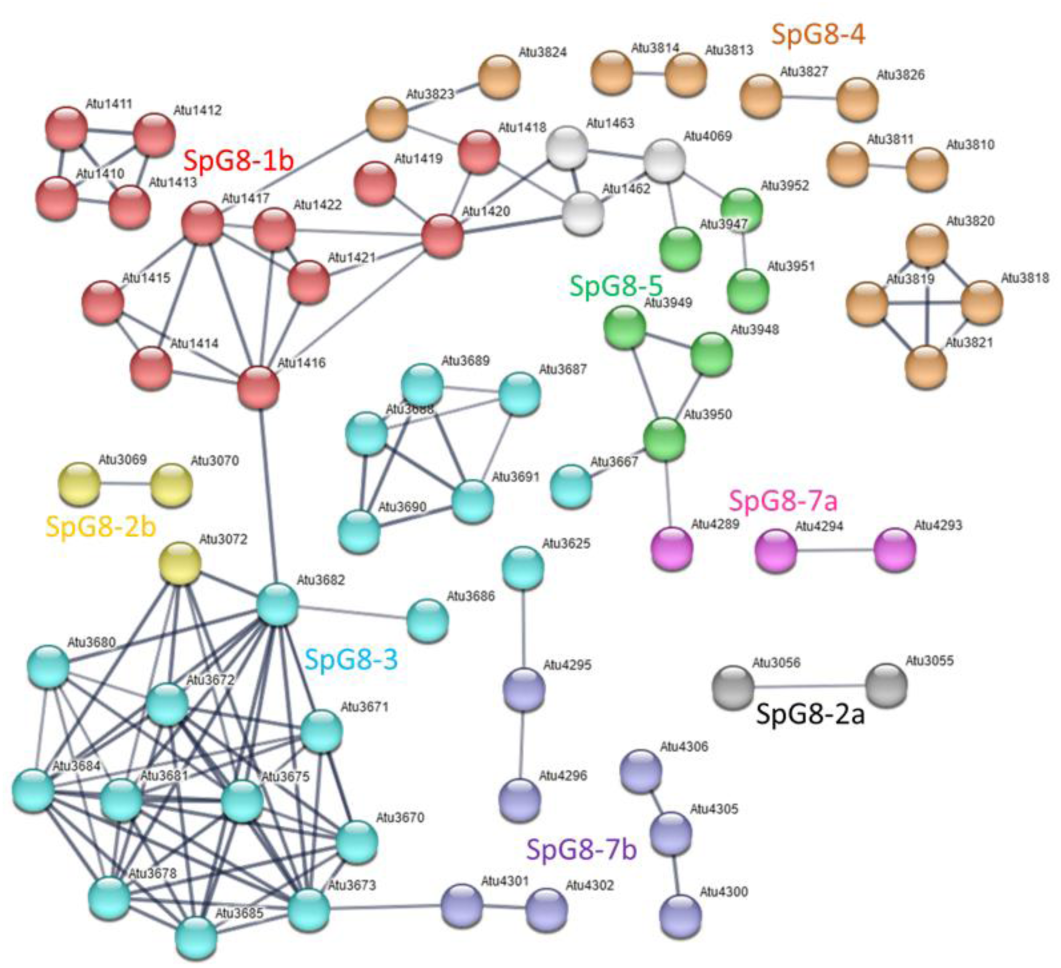
Protein–protein interaction network of the *A. fabrum* C58 species-specific proteins. The proteins are represented by nodes, whose color depends on the species-specific region they belong to; red for SpG8-1b, dark grey for SpG8-2a, yellow for SpG8-2b, cyan for SpG8-3, orange for SpG8-4, green for SpG8-5, magenta for SpG8-7a and purple for SpG8-7b. The interactions among them are represented by edges. In white, there are three proteins that do not belong to the species-specific regions but enable the connection between two of them. Proteins from species-specific regions that are not connected are not represented. The network was constructed with STRING v12, using a high confidence level of 0.7.

Mutant strains were constructed by replacing the species-specific regions with a *nptII* cassette enabling the strain to resist kanamycin/neomycin by the synthesis of neomycin phosphotransferase II protein (Lang et al., 2013). First, the non-detrimental effect of these mutations on the establishment and the survival of the mutant bacterial strains on the roots of *M. truncatula* seedlings *in vitro* cultivated was confirmed (**Fig.S2**). Secondly, root metabolite content of plants inoculated either by the wild type C58 strain (WT1) or by the kanamycin resistant C58 derivative (WT2) were compared. Only one compound was significantly affected (excluded from further analyses), showing the weak influence of neomycin phosphotransferase II protein on the plant root metabolites. Indeed, the *nptII* gene was often used as a selectable marker for the construction of GMO with no pleiotropic effects on the transcriptomes of transgenic plants (Miki et al., 2009). Moreover, these results with WT1 and WT2 underline the reproducibility of the plant reaction to the inoculation of *A. fabrum* C58 strain. Thus, in the plant experiment conducted with the C58 mutant strains, the differences in root metabolic profile observed compared to the WT condition are the result of the absence of a species-specific region rather than the presence of a resistance gene. Furthermore, the lack of effect cannot be attributed to the absence of expression *in planta* of the species-specific region (**Fig.S3**) or to the disappearance of the mutant bacterial strain on *M. truncatula* root (**Fig.S2**).

Previously, an *A. fabrum* strain (GMI9023, a C58 strain without plasmid) was reported to induce a plant-beneficial effect on maize with an increase in shoot and root dry biomass after 10 days of co-culture (Walker et al., 2013). However, after 14 days of co-culture under our conditions, aerial and root biomass obtained from *M. truncatula* were not significantly modified by inoculation with *A. fabrum* strains (wild-type C58 or deletion mutants) compared with the non-inoculated plants (**Fig. 1**). Likewise, the amounts of methanolic extract obtained from the roots were similar between all conditions (**Fig. S4**). Nevertheless, the analyses revealed a qualitative modification of these root extracts, as significant changes in the relative abundances of some compounds were highlighted between the WT condition and the NI or mutant strain conditions (**Fig. 3 and 4**). Among these highlighted discriminating metabolites, whatever the conditions, one was an amino acid, while the others were flavonoids belonging to the aurone, flavanone, isoflavone, flavone and flavonol subclasses (**Fig. 6**). The aromatic amino acid tryptophan, derived from the shikimate pathway, is a central molecule in plant metabolism. In addition to its function in protein biosynthesis, tryptophan is a precursor of a wide range of specialized metabolites and phytohormones (Tzin and Galili, 2010) and was already observed to be modulated during plant-microbe interactions (Camañes et al., 2015; Valette et al., 2019; Chen et al., 2023).

**Fig 6.**
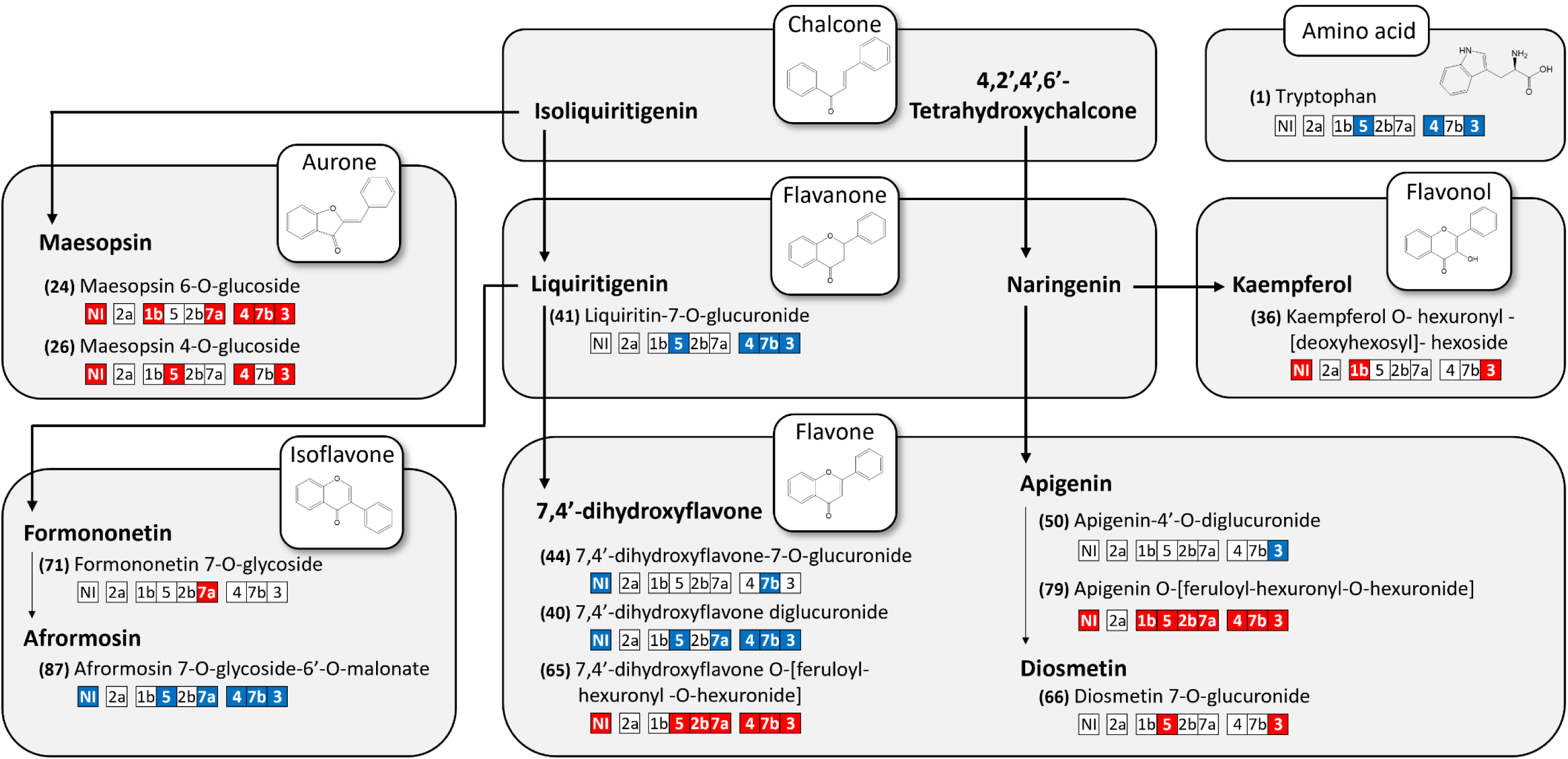
Discriminating metabolites in flavonoid biosynthesis pathways. The numbering in brackets corresponds to metabolite numbering in **Table 3** and in heatmap in Fig. 4. The discriminating compounds annotated are presented by chemical class and flavonoid subclass and plotted against their biosynthetic pathways in plants. Boxes indicate the experimental conditions of plants inoculated or not with one of the mutant strains of *A. fabrum*. Colors in boxes correspond to the overabundance (red) or underabundance (blue) of each metabolite when compared to the relative abundance of the wild-type condition. NI: non-inoculated condition, numbering 1b to 7b: plants inoculated with one of the deletion mutants of the *A. fabrum-*specific regions. 1b = C58ΔSpG8-1b; 2a = C58ΔSpG8-2a; 2b = C58ΔSpG8-2b; 3 = C58ΔSpG8-3; 4 = C58ΔSpG8-4; 5= C58ΔSpG8-5; 7a = C58ΔSpG8-7a; 7b= C58ΔSpG8-7b.

### 1. *A. fabrum* inoculation modulates plant’s flavonoid profile

When compared to uninoculated *M. truncatula* plants, the ones challenged by WT C58 strain modulated the relative abundances of some compounds among which were identified 8 flavonoids of the flavone (4), isoflavone (1), flavonol (1) and aurone (2) subclasses (**Fig. 6**). This is in line with other work highlighting the involvement of isoflavone and flavone in plant-microorganism interactions (Dixon, 1986; Schmelzer et al., 1988; Dakora and Phillips, 1996; Martens and Mithöfer, 2005; Zhang et al., 2007a). For example, isoflavones can function as allelopathic agents or as signaling molecules mediating interactions with pathogenic microorganisms or with symbiotic bacteria like *Rhizobium* (Phillips and Kapulnik, 1995; Farag et al., 2007). The isoflavone glycoside found as discriminating between the NI and the WT condition is annotated as afrormosin 7-O-glucoside-6”-O-malonate (**87**). Afrormosin, afrormosin glucoside and afrormosin glucoside malonate are the major isoflavonoids described in *M. truncatula* cell cultures (Farag et al., 2007, 2008) and have also been isolated from roots and cell suspension cultures of *M. sativa* L. (Kessmann et al., 1990). Afrormosin, might have a defensive role against both insects and fungi, but does not seem to be an antimicrobial compound (Arnoldi et al., 1986; Caballero et al., 1986; Wittstock and Gershenzon, 2002; Farag et al., 2008). The malonate conjugated forms can be used to store the less soluble isoflavone aglycones. Upon microbial infection, the aglycones are reported to be generated from the malonate conjugates (Sumner et al., 1996; Edwards et al., 1997). The inoculation by *A. fabrum* wild-type C58 strain seems to induce the increase of this molecule in roots (78% more in its relative abundance), suggesting an elicitor effect on the biosynthesis and storage of this plant defense compound.

Moreover, some flavones, including 7,4’-dihydroxyflavone, play a role as signaling molecules in interaction with other organisms. This compound is one of the most potent *nod*-gene inducers of rhizobia in *M. truncatula* (Maxwell et al., 1992; Wasson et al., 2006; Zhang et al., 2007a, 2009) since it increases in its root exudates under nitrogen starvation (Zuanazzi et al., 1998). This molecule is thus involved in a close interaction between *M. truncatula* and rhizobia that is phylogenetically closely related to *A. fabrum*. In our study, three of the flavones annotated, whose relative abundances in *M. truncatula* root were modified by the inoculation of the WT *A. fabrum* C58 strain, were hexuronide conjugates of 7,4’-dihydroxyflavone. The mono- and di-hexuronide derivatives (**40**, **44**) were enhanced in WT condition (by around 56% and 38% more in their relative abundances, respectively) while the feruloyl-dihexuronide derivative (**65**) was decreased (by around 47% less in its relative abundance), as if the feruloyl acylation had less occurred. Similarly, a feruloyl-dihexuronide derivative of apigenin (**79**, another flavone annotated) was decreased (by around 61% in its relative abundance) in the *M. truncatula* roots inoculated by WT *A. fabrum* C58 strain compared to the non-inoculated ones. However, their flavone aglycones (7,4’-dihydroxyflavone and apigenin) could be released into the plant’s root exudate when the plant perceives the WT strain, as signaling or defense molecules (Korenblum et al., 2022; Wang et al., 2022). There would therefore be less storage in feruloyl-acylated form.

More surprisingly, this metabolomic comparison revealed in the WT condition the decrease (by around 57% and 22% in their relative abundances) of two hexoside conjugates of maesopsin (**24**, **26**, respectively), which were two isomers of auronols (a small set of aurone derivatives). Among plant specialized metabolites, aurones are considered as minor flavonoids due to their low structural diversity (122 described) and limited distribution within plant biodiversity (Bohm, 1988; Boucherle et al., 2017). The maesopsin 4-O-glucoside (**26**, also named hovetrichoside C) was first described in *M. truncatula* roots in 2009 by Stochmal et al., but was already known in other plant species and was first isolated from *Hovenia trichocarea* (Yoshikawa et al., 1998), while maesopsin 6-O-glucoside (**24**) seems to have only been isolated from bark of *Ceanothus americanus* (Li et al., 1997) and stem of *Sargentodoxa cuneata* (Zeng et al., 2015). To our knowledge, these maesopsin hexoside derivatives have not been reported as metabolites involved in plant-bacteria interactions. However, other compounds belonging to aurones have been reported as phytoalexins, such as cephalocerone, the first aurone whose overproduction has been demonstrated in the cactus *Cephalocereus senilis* after microbial inoculation (Paré et al., 1991, 1992). Other aurones were reported to have antimicrobial activities against phytopathogens such as castillene D on the fungi *Lenzites trabea* (Gómez-Garibay et al., 1990) or sulfuretin on the bacterium *Ralstonia solanacearum* (Zhao et al., 2011). Furthermore, upon yeast elicitation, *M. truncatula* root cell cultures were demonstrated to overproduced two aurones, hispidol and its glucoside derivative, in the cells and with a large secretion of hispidol into the culture medium (Farag et al., 2009). Their experiments revealed the implication of some peroxidases (MtPRXs) enzymes and isoliquiritigenin as a precursor in the biosynthesis of these aurones in *M. truncatula* root cell cultures. Hispidol showed antifungal activity against *Phoma medicaginis* (a plant pathogen) and was proposed to be a potential phytoalexin (Farag et al., 2009). However, they have never detected these compounds in *M. truncatula* roots and leaves and hypothesize that these hispidols have arisen as by-products following an imbalance between the upstream and downstream segments of the phenylpropanoid pathway in this cell culture context. In our experiment, not hispidols but two other aurone hexoside derivatives (**24**, **26**) were highlighted. Their decrease in *M. truncatula* roots exposed to WT *A. fabrum* C58 strain could perhaps result from their hydrolysis and the secretion of their aglycone out of the roots in the culture medium (as defensive compounds, like hispidol) or from a limiting availability of isoliquiritigenin more redirected towards the pathways leading to the biosynthesis of certain isoflavones and flavones, like the ones (**87**, **44** and **40**) shown to be increased in parallel (**Fig.6**).

### 2. Species-specific regions involvement in the plant metabolites modifications upon *A.fabrum* inoculation

To assess the potential involvement of the species-specific gene regions of *A. fabrum* in the interaction with the plant and in the modulations of root metabolites mentioned above, root metabolite profiles of *M. truncatula* inoculated by mutant strains of these species-specific regions were compared to those inoculated by the wild-type strain C58 (WT condition). Fifteen metabolites were modulated by at least one mutant strain, nine of which were highlighted above in the *A. fabrum* - *M. truncatula* root interaction (of the eleven metabolites modulated by this interaction, **Fig.4 and 6**).

#### 2.1. The species-specific region of curdlan does not impact *M. truncatula* root phenolic compounds

In the ΔSpG8-2a condition (*i.e.* plants inoculated by the C58ΔSpG8-2a mutant strain) none of the detected root compounds had a significantly altered relative abundance when compared with the WT condition (**Fig.4**). These results suggest that this SpG8-2a specific region has no influence on these plant metabolites and would not play a direct role in the plant’s perception of the bacteria. The main predicted function of the SpG8-2a specific genes region of *A. fabrum* is curdlan biosynthesis. Curdlan is an exopolysaccharide (EPS) induced under stress conditions, such as depletion of the nitrogen source and a low pH (Harada and Harada, 1996; Kim et al., 1999; Yu et al., 2015b). This EPS surrounds bacteria, and its production could be likely to protect bacteria against environmental stresses and to increase its survival in the soil environment (Matthysse, 2018). The expression of this SpG8-2a specific-region on *M. truncatula* roots was validated with eGFP transcriptional fusion (**Fig.S3**), and, in our conditions, with or without this SpG8-2a specific-region the bacteria can similarly colonize and survive on the *M. truncatula* root system (**Fig.S2**). However, *A. fabrum* C58 strain is described to biosynthesize other EPS (Matthysse et al., 2005; Jeong et al., 2022). The lack of curdlan production by this mutant strain could be counterbalanced by another EPS produced by *A. fabrum*, like the succinoglycan. It is suggested that succinoglycan, which is produced in large amounts in this bacterium, protects *A. fabrum* from environmental stress (Matthysse, 2018). Under our experimental conditions of simple interaction of a bacterial strain with plant roots, the presence of curdlan did not appear to make a significant contribution, for either the plant or the bacteria. However, this curdlan, deposited as a capsule around *Agrobacterium* (McIntosh, 2024), may play a role in bacteria protection and could be important for its fitness in a competitive colonization of a real rhizosphere. Interestingly, the only region (SpG8-2a) that does not influence the specialized metabolites detected here in roots is also the only one whose proteins are not directly linked to those of the other specific regions (**Fig.5**).

#### 2.2. Coordinated effect of most *A. fabrum*-specific regions on phenolics in *M. truncatula* roots?

With the exception of C58ΔSpG8-2a, all mutant strains significantly altered the relative abundance of more than one phenolic compound detected in the *M. truncatula* root extracts. In addition, apart from four compounds (formonometin-7-O-glycoside (**71**), apigenin-4’-O-dihexuronide (**50**), 7,4’-dihydroxyflavone-7-O-glucuronide (**44**) and an unidentified substance (**81**)) significantly modified by a single mutant strain (C58ΔSpG8-7a, C58ΔSpG8-7b or C58ΔSpG8-3), all the discriminating root compounds were affected by the inoculation of more than one mutant strain. Among these later, derivatives of same aglycones (substituted by malonate, hexose, hexuronic acid, ferulic acid or a combination of these) were significantly modulated by the mutant strains, indicating their impact on certain metabolic pathways during interaction with *M.truncatula* roots, such as the isoflavone and flavone pathways (**Fig.6**). Furthermore, when the relative abundance of a compound was altered, it was always in the same way for all the mutant strain conditions concerned (always underabundant or overabundant relative to the WT condition), as if there was a cross-talk or coordinated effect of these species-specific regions during the interaction of *A. fabrum* with *M. truncatula* (**Fig.4 and 6**). These results are consistent with the predicted links between proteins encoded by genes in species-specific regions, where at least one protein from these regions was linked to another of them (**Fig.5**). These results suggest a scenario of ecological functioning in which *A. fabrum*-specific regions may be more closely linked than predicted by Lassalle (2011) from gene annotations.

#### 2.3 Hexuronide derivatives, an important brick in the specific niche construction of *A. fabrum* in *M. truncatula*?

Of the thirteen compounds modulated by mutant strains and annotated, twelve belong to the flavonoids, eight of which are hexuronide glycoconjugates (**Fig.4 and 6**). Among the flavonoid glycosides of plants, flavonoid hexuronides are special in that they contain hexuronic acid groups instead of or in addition to sugar units and can also be acylated with hydroxycinnamic acids (HCAs) (Kowalska et al., 2007; Marczak et al., 2010). Glycosylation and acylation have often been reported to modify the bioactivity, stability, solubility and subcellular localization of phenolic compounds. Glycosylated phenolic compounds are often regarded as storage forms for their aglycones (Bowles and Lim, 2010; Le Roy et al., 2016; Slámová et al., 2018; Johnson et al., 2021; Essa et al., 2023; Isidore et al., 2024). A wide diversity of flavonoid hexuronides were reported in roots and aerial parts of *M. truncatula* (Kowalska et al., 2007; Staszków et al., 2011; Alvarez-Rivera et al., 2022; Rai et al., 2023). Furthermore, of the mutant strains involved in the modulation of these flavonoid hexuronides, three (C58ΔSpG8-3, C58ΔSpG8-5 or C58ΔSpG8-7a) showed lower *in vitro* metabolic activity than the wild-type strain when grown on medium containing hexuronic acids (gluconic, galacturonic and glucuronic acids) (**Fig.S1**). These species-specific regions appear to influence the metabolic activity of the strain in the presence of hexuronic acids. The ability to use hexuronic acid could therefore be an important feature in the adaptation of *A. fabrum* to *M. truncatula* and in the construction of its specific niche in this rhizosphere of a plant rich in hexuronide conjugates.

#### 2.4 Feruloyl-acylated flavones influenced by the species-specific regions of *A. fabrum*

Metabolomic comparison clearly demonstrated, for almost all mutant strain conditions, the overabundance of two feruloyl-dihexuronide derivatives of flavones compared with the WT condition (**Fig.6**). Depending on the mutant strain, the feruloyl-dihexuronide derivative of 7,4’-dihydroxyflavone (**65**) and apigenin (**79**) have a relative abundance multiplied by around 1.8 to 2.2 or 2.0 to 3.7, respectively, compared with the WT condition. Whereas their mono- or di-hexuronide derivatives (**40, 44, 50**) are either less abundant or equivalent under the mutant strain conditions compared to the WT condition, as if the last step of feruloyl acylation had less occurred in plant roots for the WT condition. HCAs such as ferulic acid and its derivatives are important for *A. fabrum* ecology and fitness in either of its lifestyles pathogenic and commensal in the rhizosphere (Meyer et al., 2018, 2019). The SpG8-1b specific-region encodes the complete degradation pathway of some HCAs that will ultimately be used as a source of carbon and energy by *A. fabrum* (Campillo et al., 2014; Meyer et al., 2018). In HCAs-rich environments such as the rhizosphere (Mandal et al., 2009, 2010), *A. fabrum* could degrade and assimilate HCA to obtain a competitive advantage over other agrobacteria (Meyer et al., 2018). The importance of feruloyl derivatives during the interaction of *A. fabrum* with plants is supported by these results of modulation of feruloyl-derivatives of flavones. The underabundance of flavone feruloyl-derivatives could be linked to a release of ferulic acid or flavone aglycone into the rhizosphere when the wild-type strain is perceived by the plant, and/or to a degradation of ferulic acid by the wild-type strain only and not by the mutant strains, whatever the species-specific region. The absence of a single species-specific region has altered this influence on plant metabolism. It is already known that the SpG8-3 and the SpG8-1b region have a link between each other. Indeed, a coordinated expression has been observed following the induction with HCAs and the presence of one of them is required for the expression of the other (Baude et al., 2016). In addition, it should be noted that one of the compounds (**36**) is uniquely modulated by the absence of one of these regions SpG8-3 or SpG8-1b (**Fig.6**). This again shows the link between these species-specific regions and their impact on plant metabolites.

#### 2.5 Importance of signals in the interaction between *A. fabrum* and plant

Among the *A. fabrum*-specific regions, two of them have a greater impact on specialized plant metabolites (**Fig.4**). On the one hand, the specific region SpG8-3 appears to be somewhat central among the species-specific regions, as its proteins are suspected of interacting with proteins from four different species-specific regions (**Fig. 5**). In addition, inoculation of this mutant strain (ΔSpG8-3) had the greatest impact with modification of the highest number of root secondary metabolites (13 compounds). In contrast, proteins from the SpG8-7b specific region are suspected of being linked only to proteins from a single specific region, SpG8-3. However, inoculation of plants with the ΔSpG8-7b mutant strain alters the relative abundance of eight compounds.

The SpG8-3 region encodes the biosynthesis of a siderophore helping *A. fabrum* to grow under iron limiting conditions (Rondon et al., 2004; Vinnik et al., 2021). In the rhizosphere, the competition for iron is determinant for competitive colonization of plants. Competition (and cooperation) for iron uptake involves microorganisms but also the plant itself. An important function such as iron intake has a wide impact on the physiology of both partners in the interaction, and so has wide influence on the plant metabolism.

The predicted function of the SpG8-7b region is the involvement in the perception of environmental signals that may be responsible for the activation of other functions (Lassalle et al., 2011). Thus, this region is involved in large modification of root secondary metabolites, probably because signal detection is essential to modulate bacterial metabolism, which in turn can have an impact on plant metabolites (degradation or induction of biosynthesis). Four of the compounds modified by this ΔSpG8-7b condition belong to the same flavone biosynthesis pathway involving the 7,4’-dihydroxyflavone derivatives (**40**, **41**, **44**, **65**), one of them liquiritigenin 4’-hexosyl-7-O-hexuronide (**41**) being the precursor of the pathway (**Fig.6**). Liquiritigenin was a nod inducer gene, being then beneficial for the symbiosis legume-*Rhizobium* (Recourt et al., 1991). Moreover, as already suggested, 7,4’-dihydroxyflavone is suspected to be a signal molecule (Maxwell et al., 1992; Harrison and Dixon, 1993; Wasson et al., 2006; Zhang et al., 2007a, 2009). Thus, SpG8-7b region could be involved in the signal exchange between the bacteria and the plant. Two genes from the SpG8-7b region (Atu4300 and Atu4305) encode a two-component system and are homologous to bacterial genes whose proteins are involved in plant host recognition during the nodulation process or activate specific degradation pathways (Lau et al., 1997; Panke et al., 1998; Lang et al., 2008). Thanks to the functional similarities between homologous genes in the Rhizobiaceae, it is possible to suspect a direct role for SpG8-7b in plant recognition and thus its adaptive advantage in lifestyles that require close interactions with the plant. Given the presumed link between the SpG8-7b and SpG8-3 proteins, and the apparent pivotal role of SpG8-3, it is conceivable that SpG8-7b is dedicated to the perception of a plant signal that will be transduced and involved in the regulation of SpG8-3 expression, and through this to other specific regions of *A. fabrum*.

## CONCLUSION

Metabolomic analyses have shown that *A. fabrum* inoculation modulates the content of phenolic compounds in *M. truncatula* roots, in particular flavonoids, some of which are known to be plant defense compounds or signaling molecules. These modulations in root metabolites often appear to be linked to at least one of the *A. fabrum*-specific regions, as the relative abundance of these compounds changed upon inoculation of the mutant strains compared with the wild-type strain. Our results underline a putative cross-talk or coordinated effect of species-specific regions during this interaction, as all but one mutant condition induced similar changes on flavonoids compared to the wild-type condition (e.g. an overabundance of flavone feruloyl-dihexuronide derivatives, and an underabundance of flavone or flavanone mono- or dihexuronide derivatives). It is as if the wild-type strain caused the release or storage of aglycones or flavonoid derivatives from the plant, with the involvement of some of its *A. fabrum*-specific regions. These results suggest a model in which a compound released by the plant could be perceived through the SpG8.7b (and/or SpG8.7a) region and would be transduced and involved in the regulation of *A. fabrum*-specific regions, in particular the SpG8.3 region which appears to be central among the specific regions. These results contribute to a better understanding of the construction of the *A. fabrum* ecological niche by highlighting the importance of its specific genes in establishing this finely-tuned interaction.

## Acknowledgements

We would like to thank the CESN (Centre d’Etude des Substances Naturelles) and PARMIC (Plateforme d’Analyses des Ressources MICrobiologiques), both platforms of LEM, and their scientific and technical staffs where this work was performed. We also thank Tony Campillo for its skillful technical assistance in setting up plant cultivation and inoculation systems and for the harvesting of samples; David Chapulliot and Sébastien Renoud for their technical assistance for the plant harvesting; Théo Pottier for his technical assistance in obtaining the control bacterial extract. We thank Elise Lacroix for some plant cultures at the “Serre et Chambres Climatiques” platform (UCBL1). Finally, we gratefully thank Pr. Laurence Voutquenne-Nazabadioko from the Reims Institute of Molecular Chemistry (ICMR, UMR CNRS 7312) for kindly provided a sample of purified Maesopsin 4-O-glycoside, and Dr. Christine Lesignor (Research Engineer at INRAE, center of Dijon) from the UMR1347 Agroecologie, GEAPSI - Equipe FILEAS, for providing us with seeds of *Medicago truncatula* A17 (Jemalong).

## Author contributions

CL, LV, and IK designed the research and performed the plant inoculation experiments; RP and LV realised the mutant strains and transcriptional fusion constructions and their analyses; RP and FD performed the root colonization experiments; IK performed root metabolites extractions, their analyses, and data curation; VG participated to the methodology and root extracts analyses; RP, THTN, GC, GM and IK performed the spectral data interpretations and the metabolite identifications; RP, THTN, LV, CL and IK performed experimental data analyses; RP, THTN and IK performed statistical analyses; RP, LV, CL and IK prepared tables and figures; RP, LV, XN, CL and IK wrote the original draft manuscript; LV, GC, CL and IK proof-read and edit the manuscript; XN, LV, CL and IK performed funding acquisition and research supervision.

## References

Al Arni, S., Drake, A.F., Del Borghi, M., and Converti, A. (2010). Study of aromatic compounds derived from sugarcane bagasse. Part I: Effect of ph. Chem. Eng. Technol. 33: 895–901.

Allaway, D., Schofield, N.A., Leonard, M.E., Gilardoni, L., Finan, T.M., and Poole, P.S. (2001). Use of differential fluorescence induction and optical trapping to isolate environmentally induced genes. Environ. Microbiol. 3: 397–406.

Alvarez-Rivera, G., Sanz, A., Cifuentes, A., Ibánez, E., Paape, T., Lucas, M.M., and Pueyo, J.J. (2022). Flavonoid accumulation varies in *Medicago truncatula* in response to mercury stress. Front. Plant Sci. 13: 933209.

Arnoldi, A., Farina, G., Galli, R., Merlini, L., and Parrino, M.G. (1986). Analogs of phytoalexins. Synthesis of some 3-phenylcoumarins and their fungicidal activity. J. Agric. Food Chem. 34: 185–188.

Badri, D.V., Weir, T.L., van der Lelie, D., and Vivanco, J.M. (2009). Rhizosphere chemical dialogues: plant-microbe interactions. Curr. Opin. Biotechnol. 20: 642–650.

Baude, J., Vial, L., Villard, C., Campillo, T., Lavire, C., Nesme, X., and Hommais, F. (2016). Coordinated regulation of species-specific hydroxycinnamic acid degradation and siderophore biosynthesis pathways in *Agrobacterium fabrum*. Appl. Environ. Microbiol. 82: 3515–3524.

Beninger, C.W. and Hall, J.C. (2005). Allelopathic activity of luteolin 7-*O*-β-glucuronide isolated from *Chrysanthemum morifolium* L. Biochem. Syst. Ecol. 33: 103–111.

Bennett, R.N. and Wallsgrove, R.M. (1994). Secondary metabolites in plant defence mechanisms. New Phytol. 127: 617–633.

Bohm, B.A. (1988). The minor flavonoids. In The Flavonoids (Springer, Boston, MA), pp. 329–388.

Boucherle, B., Peuchmaur, M., Boumendjel, A., and Haudecoeur, R. (2017). Occurrences, biosynthesis and properties of aurones as high-end evolutionary products. Phytochemistry 142: 92–111.

Bouri, M., Chattaoui, M., Gharsa, H.B., McClean, A., Kluepfel, D., Nesme, X., and Rhouma, A. (2016). Analysis of *Agrobacterium* populations isolated from Tunisian soils: genetic structure, avirulent-virulent ratios and characterization of tumorigenic isolates. J. Plant Pathol. 98: 265–274.

Bowles, D. and Lim, E.-K. (2010). Glycosyltransferases of small molecules: their roles in plant biology. In eLS (John Wiley & Sons, Ltd).

Budzianowski, J. (1991). Six flavonol glucuronides from *Tulipa gesneriana*. Phytochemistry 30: 1679–1682.

Caballero, P., Smith, C.M., Fronczek, F.R., and Fischer, N.H. (1986). Isoflavones from an insect-resistant variety of soybean and the molecular structure of afrormosin. J. Nat. Prod. 49: 1126–1129.

Camañes, G., Scalschi, L., Vicedo, B., González-Bosch, C., and García-Agustín, P. (2015). An untargeted global metabolomic analysis reveals the biochemical changes underlying basal resistance and priming in *Solanum lycopersicum*, and identifies 1-methyltryptophan as a metabolite involved in plant responses to *Botrytis cinerea* and *Pseudomonas syringae*. Plant J. 84: 125–139.

Campillo, T., Renoud, S., Kerzaon, I., Vial, L., Baude, J., Gaillard, V., Bellvert, F., Chamignon, C., Comte, G., Nesme, X., Lavire, C., and Hommais, F. (2014). Analysis of hydroxycinnamic acid degradation in *Agrobacterium fabrum* reveals a coenzyme A-dependent, beta-oxidative deacetylation pathway. Appl. Environ. Microbiol. 80: 3341–3349.

Chen, L., Ma, Y., He, T., Chen, T., Pan, Y., Zhou, D., Li, X., Lu, Y., Wu, Q., and Wang, L. (2023). Integrated transcriptome and metabolome analysis unveil the response mechanism in wild rice (*Zizania latifolia* griseb.) against sheath rot infection. Front. Genet. 14: 1163464.

Chilton, M.-D., Currier, T.C., Farrand, S.K., Bendich, A.J., Gordon, M.P., and Nester, E.W. (1974). *Agrobacterium tumefaciens* DNA and PS8 bacteriophage DNA not detected in crown gall tumors. Proc. Natl. Acad. Sci. U. S. A. 71: 3672–3676.

Cook, D.R. (1999). *Medicago truncatula* - a model in the making! Curr. Opin. Plant Biol. 2: 301–304.

Costechareyre, D., Rhouma, A., Lavire, C., Portier, P., Chapulliot, D., Bertolla, F., Boubaker, A., Dessaux, Y., and Nesme, X. (2010). Rapid and efficient identification of *Agrobacterium* species by recA allele analysis: *Agrobacterium* recA diversity. Microb. Ecol. 60: 862–872.

Dakora, F.D. and Phillips, D.A. (1996). Diverse functions of isoflavonoids in legumes transcend anti-microbial definitions of phytoalexins. Physiol. Mol. Plant Pathol. 49: 1–20.

Datsenko, K.A. and Wanner, B.L. (2000). One-step inactivation of chromosomal genes in *Escherichia coli* K-12 using PCR products. Proc. Natl. Acad. Sci. U. S. A. 97: 6640–6645.

Dessaux, Y. and Faure, D. (2018). Niche construction and exploitation by *Agrobacterium*: how to survive and face competition in soil and plant habitats. Curr. Top. Microbiol. Immunol. 418: 55–86.

Dixon, R.A. (1986). The phytoalexin response: elicitation, signalling and control of host gene expression. Biol. Rev. 61: 239–291.

Du, Y., Zou, J., Yin, Z., and Chen, T. (2023). Pan-chromosome and comparative analysis of *Agrobacterium fabrum* reveal important traits concerning the genetic diversity, evolutionary dynamics, and niche adaptation of the species. Microbiol. Spectr. 11: e0292422.

Edwards, R., Tiller, S.A., and Parry, A.D. (1997). The effect of plant age and nodulation on the isoflavonoid content of red clover (*Trifolium pratense*). J. Plant Physiol. 150: 603–610.

Essa, A.F., Teleb, M., El-Kersh, D.M., El Gendy, A.E.-N.G., Elshamy, A.I., and Farag, M.A. (2023). Natural acylated flavonoids: their chemistry and biological merits in context to molecular docking studies. Phytochem. Rev. 22: 1469–1508.

Farag, M.A., Deavours, B.E., de Fátima, A., Naoumkina, M., Dixon, R.A., and Sumner, L.W. (2009). Integrated metabolite and transcript profiling identify a biosynthetic mechanism for hispidol in *Medicago truncatula* cell cultures. Plant Physiol. 151: 1096–1113.

Farag, M.A., Huhman, D.V., Dixon, R.A., and Sumner, L.W. (2008). Metabolomics reveals novel pathways and differential mechanistic and elicitor-specific responses in phenylpropanoid and isoflavonoid biosynthesis in *Medicago truncatula* cell cultures. Plant Physiol. 146: 387–402.

Farag, M.A., Huhman, D.V., Lei, Z., and Sumner, L.W. (2007). Metabolic profiling and systematic identification of flavonoids and isoflavonoids in roots and cell suspension cultures of *Medicago truncatula* using HPLC-UV-ESI-MS and GC-MS. Phytochemistry 68: 342–354.

Ferreres, F., Grosso, C., Gil-Izquierdo, A., Valentão, P., Azevedo, C., and Andrade, P.B. (2014). HPLC-DAD-ESI/MSn analysis of phenolic compounds for quality control of *Grindelia robusta* Nutt. and bioactivities. J. Pharm. Biomed. Anal. 94: 163–172.

Fletcher, J.S., Kotze, H.L., Armitage, E.G., Lockyer, N.P., and Vickerman, J.C. (2013). Evaluating the challenges associated with time-of-fight secondary ion mass spectrometry for metabolomics using pure and mixed metabolites. Metabolomics 9: 535–544.

Fu, F. and Wang, H.L. (2015). Metabolomics reveals consistency of the shoot system in *Medicago truncatula* by HPLC-UV-ESI-MS/MS. Int. J. Food Sci. Technol. 50: 2183–2192.

Gómez-Garibay, F., Chilpa, R.R., Quijano, L., Calderón Pardo, JoséS., and Ríos Castillo, T. (1990). Methoxy furan auranols with fungistatic activity from *Lonchocarpus castilloi*. Phytochemistry 29: 459–463.

González-Mula, A., Lang, J., Grandclément, C., Naquin, D., Ahmar, M., Soulère, L., Queneau, Y., Dessaux, Y., and Faure, D. (2018). Lifestyle of the biotroph *Agrobacterium tumefaciens* in the ecological niche constructed on its host plant. New Phytol. 219: 350–362.

Greenham, J., Harborne, J.B., and Williams, C.A. (2003). Identification of lipophilic flavones and flavonols by comparative HPLC, TLC and UV spectral analysis. Phytochem. Anal. PCA 14: 100–118.

Gupta, S., Schillaci, M., and Roessner, U. (2022). Metabolomics as an emerging tool to study plant–microbe interactions. Emerg. Top. Life Sci. 6: 175–183.

Guy, P.A., Renouf, M., Barron, D., Cavin, C., Dionisi, F., Kochhar, S., Rezzi, S., Williamson, G., and Steiling, H. (2009). Quantitative analysis of plasma caffeic and ferulic acid equivalents by liquid chromatography tandem mass spectrometry. J. Chromatogr. B Analyt. Technol. Biomed. Life. Sci. 877: 3965–3974.

Harada, T. and Harada, A. (1996). Curdlan and succinoglycan In Dimitrio, S.

Harrison, M.J. and Dixon, R.A. (1993). Isoflavonoid accumulation and expression of defense gene transcripts during the establishment of vesicular-arbuscular mycorrhizal associations in roots of *Medicago truncatula*. Mol. Plant. Microbe Interact. 6: 643.

Hooykaas, P.J.J. (2023). The Ti plasmid, driver of *Agrobacterium* pathogenesis. Phytopathology 113: 594–604.

Ibrahim, A.R. and Abul-Hajj, Y.J. (1990). Aromatase inhibition by flavonoids. J. Steroid Biochem. Mol. Biol. 37: 257–260.

Isidore, E., Willig, G., Brunissen, F., Magro, C., Monteux, C., and Ioannou, I. (2024). Selective recovery of glycosylated phenolic compounds from nectarine tree branches (*Prunus persica* var. *nucipersica*). Food Chem. Adv. 4: 100585.

Jan, R., Asaf, S., Numan, M., Lubna, and Kim, K.-M. (2021). Plant secondary metabolite biosynthesis and transcriptional regulation in response to biotic and abiotic stress conditions. Agronomy 11: 968.

Jasiński, M., Kachlicki, P., Rodziewicz, P., Figlerowicz, M., and Stobiecki, M. (2009). Changes in the profile of flavonoid accumulation in *Medicago truncatula* leaves during infection with fungal pathogen *Phoma medicaginis*. Plant Physiol. Biochem. 47: 847–853.

Jeong, J.-P., Kim, Y., Hu, Y., and Jung, S. (2022). Bacterial succinoglycans: structure, physical properties, and applications. Polymers 14: 276.

Jiang, P., Dai, W., Yan, S., Chen, Z., Xu, R., Ding, J., Xiang, L., Wang, S., Liu, R., and Zhang, W. (2011). Potential biomarkers in the urine of myocardial infarction rats: a metabolomic method and its application. Mol. Biosyst. 7: 824–831.

Johnson, J.B., Mani, J.S., Broszczak, D., Prasad, S.S., Ekanayake, C.P., Strappe, P., Valeris, P., and Naiker, M. (2021). Hitting the sweet spot: A systematic review of the bioactivity and health benefits of phenolic glycosides from medicinally used plants. Phytother. Res. 35: 3484–3508.

Jung, J.-Y., Jung, Y., Kim, J.-S., Ryu, D.H., and Hwang, G.-S. (2013). Assessment of peeling of *Astragalus* roots using 1H NMR- and UPLC-MS-based metabolite profiling. J. Agric. Food Chem. 61: 10398–10407.

Kessmann, H., Edwards, R., Geno, P.W., and Dixon, R.A. (1990). Stress responses in alfalfa (*Medicago sativa* L.): v. constitutive and elicitor-induced accumulation of isoflavonoid conjugates in cell suspension cultures. Plant Physiol. 94: 227–232.

Kim, M.-K., Lee, I.-Y., Ko, J.-H., Rhee, Y.-H., and Park, Y.-H. (1999). Higher intracellular levels of uridinemonophosphate under nitrogen-limited conditions enhance metabolic flux of curdlan synthesis in *Agrobacterium* species. Biotechnol. Bioeng. 62: 317–323.

Koprivova, A. and Kopriva, S. (2022). Plant secondary metabolites altering root microbiome composition and function. Curr. Opin. Plant Biol. 67: 102227.

Korenblum, E., Massalha, H., and Aharoni, A. (2022). Plant–microbe interactions in the rhizosphere via a circular metabolic economy. Plant Cell 34: 3168–3182.

Kowalska, I., Stochmal, A., Kapusta, I., Janda, B., Pizza, C., Piacente, S., and Oleszek, W. (2007). Flavonoids from barrel medic (*Medicago truncatula*) aerial parts. J. Agric. Food Chem. 55: 2645–2652.

Lang, J., Planamente, S., Mondy, S., Dessaux, Y., Moréra, S., and Faure, D. (2013). Concerted transfer of the virulence Ti plasmid and companion At plasmid in the *Agrobacterium tumefaciens*-induced plant tumour. Mol. Microbiol. 90: 1178–1189.

Lang, K., Lindemann, A., Hauser, F., and Göttfert, M. (2008). The genistein stimulon of *Bradyrhizobium japonicum*. Mol. Genet. Genomics MGG 279: 203–211.

Lassalle, F. et al. (2017). Ancestral genome estimation reveals the history of ecological diversification in *Agrobacterium*. Genome Biol. Evol. 9: 3413–3431.

Lassalle, F. et al. (2011). Genomic species are ecological species as revealed by comparative genomics in *Agrobacterium tumefaciens*. Genome Biol. Evol. 3: 762–781.

Lau, P.C., Wang, Y., Patel, A., Labbé, D., Bergeron, H., Brousseau, R., Konishi, Y., and Rawlings, M. (1997). A bacterial basic region leucine zipper histidine kinase regulating toluene degradation. Proc. Natl. Acad. Sci. U. S. A. 94: 1453–1458.

Le Roy, J., Huss, B., Creach, A., Hawkins, S., and Neutelings, G. (2016). Glycosylation is a major regulator of phenylpropanoid availability and biological activity in plants. Front. Plant Sci. 7: 735.

Li, Q., Liu, Y., Han, L., Liu, J., Liu, W., Feng, F., Zhang, J., and Xie, N. (2016). Chemical constituents and quality control of two *Dracocephalum* species based on high-performance liquid chromatographic fingerprints coupled with tandem mass spectrometry and chemometrics. J. Sep. Sci. 39: 4071–4085.

Li, X.-C., Cai, L., and D. Wu, C. (1997). Antimicrobial compounds from *Ceanothus americanus* against oral pathogens. Phytochemistry 46: 97–102.

Ma, H., Liu, Y., Mai, X., Liao, Y., Zhang, K., Liu, B., Xie, X., and Du, Q. (2016). Identification of the constituents and metabolites in rat plasma after oral administration of HuanglianShangqing pills by ultra high-performance liquid chromatography/quadrupole time-of-flight mass spectrometry. J. Pharm. Biomed. Anal. 125: 194–204.

Mabry, T., Markham, K.R., and Thomas, M.B. (1970). The Systematic Identification of Flavonoids (Springer-Verlag: Berlin Heidelberg).

Mandal, S., Mitra, A., and Mallick, N. (2009). Time course study on accumulation of cell wall-bound phenolics and activities of defense enzymes in tomato roots in relation to *Fusarium* wilt. World J. Microbiol. Biotechnol. 25: 795–802.

Mandal, S.M., Chakraborty, D., and Dey, S. (2010). Phenolic acids act as signaling molecules in plant-microbe symbioses. Plant Signal. Behav. 5: 359–368.

Marczak, Ł., Stobiecki, M., Jasiński, M., Oleszek, W., and Kachlicki, P. (2010). Fragmentation pathways of acylated flavonoid diglucuronides from leaves of *Medicago truncatula*. Phytochem. Anal. 21: 224–233.

Marczak, Ł., Znajdek-Awiżeń, P., and Bylka, W. (2016). The use of mass spectrometric techniques to differentiate isobaric and isomeric flavonoid conjugates from *Axyris amaranthoides*. Mol. Basel Switz. 21: 1229.

Martens, S. and Mithöfer, A. (2005). Flavones and flavone synthases. Phytochemistry 66: 2399–2407.

Massalha, H., Korenblum, E., Tholl, D., and Aharoni, A. (2017). Small molecules below-ground: the role of specialized metabolites in the rhizosphere. Plant J. Cell Mol. Biol. 90: 788–807.

Matthysse, A.G. (2018). Exopolysaccharides of *Agrobacterium tumefaciens*. In Agrobacterium Biology: From Basic Science to Biotechnology, S.B. Gelvin, ed, Current Topics in Microbiology and Immunology. (Springer International Publishing: Cham), pp. 111–141.

Matthysse, A.G., Marry, M., Krall, L., Kaye, M., Ramey, B.E., Fuqua, C., and White, A.R. (2005). The effect of cellulose overproduction on binding and biofilm formation on roots by *Agrobacterium tumefaciens*. Mol. Plant-Microbe Interactions® 18: 1002–1010.

Maxwell, C.A., Edwards, R., and Dixon, R.A. (1992). Identification, purification, and characterization of *S*-adenosyl-l-methionine: Isoliquiritigenin 2′-*O*-methyltransferase from alfalfa (*Medicago sativa* L.). Arch. Biochem. Biophys. 293: 158–166.

McIntosh, M. (2024). Genetic engineering of *Agrobacterium* increases curdlan production through increased expression of the crdASC genes. Microorganisms 12: 55.

McIntosh, M., Stone, B.A., and Stanisich, V.A. (2005). Curdlan and other bacterial (1-->3)-beta-D-glucans. Appl. Microbiol. Biotechnol. 68: 163–173.

Meyer, T., Renoud, S., Vigouroux, A., Miomandre, A., Gaillard, V., Kerzaon, I., Prigent-Combaret, C., Comte, G., Moréra, S., Vial, L., and Lavire, C. (2018). Regulation of hydroxycinnamic acid degradation drives *Agrobacterium fabrum* lifestyles. Mol. Plant. Microbe Interact. 31: 814–822.

Meyer, T., Thiour-Mauprivez, C., Wisniewski-Dyé, F., Kerzaon, I., Comte, G., Vial, L., and Lavire, C. (2019). Ecological conditions and molecular determinants involved in *Agrobacterium* lifestyle in tumors. Front. Plant Sci. 10: 978.

Miki, B., Abdeen, A., Manabe, Y., and MacDonald, P. (2009). Selectable marker genes and unintended changes to the plant transcriptome. Plant Biotechnol. J. 7: 211–218.

Miotto-Vilanova, L., Courteaux, B., Padilla, R., Rabenoelina, F., Jacquard, C., Clément, C., Comte, G., Lavire, C., Ait Barka, E., Kerzaon, I., and Sanchez, L. (2019). Impact of *Paraburkholderia phytofirmans* PsJN on grapevine phenolic metabolism. Int. J. Mol. Sci. 20: 5775–5807.

Muhammad, D., Lalun, N., Bobichon, H., Le Magrex Debar, E., Gangloff, S.C., Nour, M., and Voutquenne-Nazabadioko, L. (2017). Triterpenoid saponins and other glycosides from the stems and bark of *Jaffrea xerocarpa* and their biological activity. Phytochemistry 141: 121–130.

Naqqash, T., Hameed, S., Imran, A., Hanif, M.K., Majeed, A., and van Elsas, J.D. (2016). Differential response of potato toward inoculation with taxonomically diverse plant growth promoting Rhizobacteria. Front. Plant Sci. 7: 144.

Nesme, X., Michel, M.F., and Digat, B. (1987). Population heterogeneity of *Agrobacterium tumefaciens* in galls of *Populus* L. from a single nursery. Appl. Environ. Microbiol. 53: 655–659.

Nong, Q., Malviya, M.K., Solanki, M.K., Lin, L., Xie, J., Mo, Z., Wang, Z., Song, X., Huang, X., Li, C., and Li, Y. (2023). Integrated metabolomic and transcriptomic study unveils the gene regulatory mechanisms of sugarcane growth promotion during interaction with an endophytic nitrogen-fixing bacteria. BMC Plant Biol. 23: 54.

Panke, S., Witholt, B., Schmid, A., and Wubbolts, M.G. (1998). Towards a biocatalyst for (S)-styrene oxide production: characterization of the styrene degradation pathway of *Pseudomonas* sp. strain VLB120. Appl. Environ. Microbiol. 64: 2032–2043.

Paré, P.W., Dmitrieva, N., and Mabry, T.J. (1991). Phytoalexin aurone induced in *Cephalocereus senilis* liquid suspension culture. Phytochemistry 30: 1133–1135.

Paré, P.W., Mischke, C.F., Edwards, R., Dixon, R.A., Norman, H.A., and Mabry, T.J. (1992). Induction of phenylpropanoid pathway enzymes in elicitor-treated cultures of*Cephalocereus senilis*. Phytochemistry 31: 149–153.

Park, J.A., Kim, H.J., Jin, C., Lee, K.-T., and Lee, Y.S. (2003). A new pterocarpan, (-)-maackiain sulfate, from the roots of *Sophora subprostrata*. Arch. Pharm. Res. 26: 1009–1013.

Parke, D., Ornston, L.N., and Nester, E.W. (1987). Chemotaxis to plant phenolic inducers of virulence genes is constitutively expressed in the absence of the Ti plasmid in *Agrobacterium tumefaciens*. J. Bacteriol. 169: 5336–5338.

Philippot, L., Raaijmakers, J.M., Lemanceau, P., and van der Putten, W.H. (2013). Going back to the roots: the microbial ecology of the rhizosphere. Nat. Rev. Microbiol. 11: 789–799.

Phillips, D.A. and Kapulnik, Y. (1995). Plant isoflavonoids, pathogens and symbionts. Trends Microbiol. 3: 58–64.

Pothier, J.F., Wisniewski-Dyé, F., Weiss-Gayet, M., Moënne-Loccoz, Y., and Prigent-Combaret, C. (2007). Promoter-trap identification of wheat seed extract-induced genes in the plant-growth-promoting rhizobacterium *Azospirillum brasilense* Sp245. Microbiol. Read. Engl. 153: 3608–3622.

Qiao, X., Ye, M., Xiang, C., Wang, Q., Liu, C.-F., Miao, W.-J., and Guo, D.-A. (2012). Analytical strategy to reveal the *in vivo* process of multi-component herbal medicine: a pharmacokinetic study of licorice using liquid chromatography coupled with triple quadrupole mass spectrometry. J. Chromatogr. A 1258: 84–93.

Quandt, J. and Hynes, M.F. (1993). Versatile suicide vectors which allow direct selection for gene replacement in gram-negative bacteria. Gene 127: 15–21.

Rai, N., Neugart, S., Schröter, D., Lindfors, A.V., and Aphalo, P.J. (2023). Responses of flavonoids to solar UV radiation and gradual soil drying in two *Medicago truncatula* accessions. Photochem. Photobiol. Sci. Off. J. Eur. Photochem. Assoc. Eur. Soc. Photobiol. 22: 1637–1654.

Recourt, K., Schripsema, J., Kijne, J.W., van Brussel, A.A., and Lugtenberg, B.J. (1991). Inoculation of *Vicia sativa* subsp. *nigr*a roots with *Rhizobium leguminosarum* biovar *viciae* results in release of nod gene activating flavanones and chalcones. Plant Mol. Biol. 16: 841–852.

Rondon, M.R., Ballering, K.S., and Thomas, M.G. (2004). Identification and analysis of a siderophore biosynthetic gene cluster from *Agrobacterium tumefaciens* C58. Microbiology 150: 3857–3866.

Saleh, N.A.M., Boulos, L., El-Negoumy, S.I., and Abdalla, M.F. (1982). A comparative study of the flavonoids of *Medicago radiata* with other *Medicago* and related *Trigonella* species. Biochem. Syst. Ecol. 10: 33–36.

Schmelzer, E., Jahnen, W., and Hahlbrock, K. (1988). *In situ* localization of light-induced chalcone synthase mRNA, chalcone synthase, and flavonoid end products in epidermal cells of parsley leaves. Proc. Natl. Acad. Sci. U. S. A. 85: 2989–2993.

Shams, M., Campillo, T., Lavire, C., Muller, D., Nesme, X., and Vial, L. (2012). Rapid and efficient methods to isolate, type strains and determine species of *Agrobacterium* spp. in pure culture and complex environments. Biochem. Test.

Singh, G., Agrawal, H., and Bednarek, P. (2023). Specialized metabolites as versatile tools in shaping plant– microbe associations. Mol. Plant 16: 122–144.

Singh, R., Wu, B., Tang, L., Liu, Z., and Hu, M. (2010). Identification of the position of mono-O-glucuronide of flavones and flavonols by analyzing shift in online UV spectrum (lambda max) generated from an online diode array detector. J. Agric. Food Chem. 58: 9384–9395.

Slámová, K., Kapešová, J., and Valentová, K. (2018). “Sweet flavonoids”: glycosidase-catalyzed modifications. Int. J. Mol. Sci. 19: 2126.

Staszków, A., Swarcewicz, B., Banasiak, J., Muth, D., Jasiński, M., and Stobiecki, M. (2011). LC/MS profiling of flavonoid glycoconjugates isolated from hairy roots, suspension root cell cultures and seedling roots of *Medicago truncatula*. Metabolomics 7: 604–613.

Stochmal, A., Kowalska, I., Janda, B., Perrone, A., Piacente, S., and Oleszek, W. (2009). Gentisic acid conjugates of *Medicago truncatula* roots. Phytochemistry 70: 1272–1276.

Stochmal, A., Piacente, S., Pizza, C., De Riccardis, F., Leitz, R., and Oleszek, W. (2001a). Alfalfa (*Medicago sativa* L.) flavonoids. 1. Apigenin and luteolin glycosides from aerial parts. J. Agric. Food Chem. 49: 753–758.

Stochmal, A., Simonet, A.M., Macias, F.A., Oliveira, M.A., Abreu, J.M., Nash, R., and Oleszek, W. (2001b). Acylated apigenin glycosides from alfalfa (*Medicago sativa* L.) var. Artal. Phytochemistry 57: 1223–1226.

Sumner, L.W., Paiva, N.L., Dixon, R.A., and Geno, P.W. (1996). High-performance liquid chromatography/continuous-flow liquid secondary ion mass spectrometry of flavonoid glycosides in leguminous plant extracts. J. Mass Spectrom. 31: 472–485.

Szklarczyk, D. et al. (2023). The STRING database in 2023: protein-protein association networks and functional enrichment analyses for any sequenced genome of interest. Nucleic Acids Res. 51: D638–D646.

Thuy, T.T., Kamperdick, C., Ninh, P.T., Lien, T.P., Thao, T.T.P., and Sung, T.V. (2004). Immunosuppressive auronol glycosides from *Artocarpus tonkinensis*. Pharm. 59: 297–300.

Thuy, T.T., Thien, D.D., Quang Hung, T., Tam, N.T., Anh, N.T.H., Nga, N.T., Cuc, N.T., Mai, L.P., Van Sung, T., Delfino, D.V., and Thao, D.T. (2016). *In vivo* anticancer activity of maesopsin 4-*O*-*β*-glucoside isolated from leaves of *Artocarpus tonkinensis* A. Chev. Ex Gagnep. Asian Pac. J. Trop. Med. 9: 351–356.

Tibe, O., Meagher, L.P., Fraser, K., and Harding, D.R.K. (2011). Condensed tannins and flavonoids from the forage legume sulla (*Hedysarum coronarium*). J. Agric. Food Chem. 59: 9402–9409.

Tzin, V. and Galili, G. (2010). The biosynthetic pathways for shikimate and aromatic amino acids in *Arabidopsis thaliana*. Arab. Book 2010: e0132.

Valette, M., Rey, M., Gerin, F., Comte, G., and Wisniewski-Dyé, F. (2019). A common metabolomic signature is observed upon inoculation of rice roots with various rhizobacteria. J. Integr. Plant Biol. 62: 228–246.

Vinnik, V., Zhang, F., Park, H., Cook, T.B., Throckmorton, K., Pfleger, B.F., Bugni, T.S., and Thomas, M.G. (2021). Structural and biosynthetic analysis of the fabrubactins, unusual siderophores from *Agrobacterium fabrum* strain C58. ACS Chem. Biol. 16: 125–135.

Vogel, J., Normand, P., Thioulouse, J., Nesme, X., and Grundmann, G.L. (2003). Relationship between spatial and genetic distance in *Agrobacterium* spp. in 1 cubic centimeter of soil. Appl. Environ. Microbiol. 69: 1482– 1487.

Walker, V., Bertrand, C., Bellvert, F., Moënne-Loccoz, Y., Bally, R., and Comte, G. (2011). Host plant secondary metabolite profiling shows a complex, strain-dependent response of maize to plant growth-promoting rhizobacteria of the genus *Azospirillum*. New Phytol. 189: 494–506.

Walker, V., Bruto, M., Bellvert, F., Bally, R., Muller, D., Prigent-Combaret, C., Moënne-Loccoz, Y., and Comte, G. (2013). Unexpected phytostimulatory behavior for *Escherichia coli* and *Agrobacterium tumefaciens* model strains. Mol. Plant-Microbe Interact. MPMI 26: 495–502.

Wang, L., Chen, M., Lam, P.-Y., Dini-Andreote, F., Dai, L., and Wei, Z. (2022). Multifaceted roles of flavonoids mediating plant-microbe interactions. Microbiome 10: 233.

Wang, X., Zhang, J., Lu, X., Bai, Y., and Wang, G. (2023). Two diversities meet in the rhizosphere: root specialized metabolites and microbiome. J. Genet. Genomics.

Wasson, A.P., Pellerone, F.I., and Mathesius, U. (2006). Silencing the flavonoid pathway in *Medicago truncatula* inhibits root nodule formation and prevents auxin transport regulation by rhizobia. Plant Cell 18: 1617–1629.

Wen, X.-D., Qi, L.-W., Chen, J., Song, Y., Yi, L., Yang, X.-W., and Li, P. (2007). Analysis of interaction property of bioactive components in Danggui Buxue Decoction with protein by microdialysis coupled with HPLC–DAD–MS. J. Chromatogr. B 852: 598–604.

Whitehead, D.C., Dibb, H., and Hartley, R.D. (1983). Bound phenolic compounds in water extracts of soils, plant roots and leaf litter. Soil Biol. Biochem. 15: 133–136.

Wittstock, U. and Gershenzon, J. (2002). Constitutive plant toxins and their role in defense against herbivores and pathogens. Curr. Opin. Plant Biol. 5: 300–307.

Wong, Y.C., Zhang, L., Lin, G., and Zuo, Z. (2009). Intestinal first-pass glucuronidation activities of selected dihydroxyflavones. Int. J. Pharm. 366: 14–20.

Wu, Q., Wang, M., and Simon, J.E. (2003). Determination of isoflavones in red clover and related species by high-performance liquid chromatography combined with ultraviolet and mass spectrometric detection. J. Chromatogr. A 1016: 195–209.

Ye, X., Tang, M., Chen, L., Peng, A., Ma, L., and Ye, H. (2009). Rapid separation and identification of major constituents in *Pseudolarix kaempferi* by ultra-performance liquid chromatography coupled with electrospray and quadrupole time-of-flight tandem mass spectrometry. Rapid Commun. Mass Spectrom. 23: 3954–3962.

Yoshikawa, K., Kondo, Y., Kimura, E., and Arihara, S. (1998). A lupane-triterpene and a 3(2→1)abeolupane glucoside from *Hovenia trichocarea*. Phytochemistry 49: 2057–2060.

Yu, N., He, C., Awuti, G., Zeng, C., Xing, J., and Huang, W. (2015a). Simultaneous determination of six active compounds in Yixin Badiranjibuya granules, a traditional chinese medicine, by RP-HPLC-UV method. J. Anal. Methods Chem. 2015: 974039.

Yu, X., Zhang, C., Yang, L., Zhao, L., Lin, C., Liu, Z., and Mao, Z. (2015b). CrdR function in a curdlan-producing *Agrobacterium* sp. ATCC31749 strain. BMC Microbiol. 15: 25.

Zeiss, D.R., Mhlongo, M.I., Tugizimana, F., Steenkamp, P.A., and Dubery, I.A. (2019). Metabolomic profiling of the host response of tomato (*Solanum lycopersicum*) following infection by *Ralstonia solanacearum*. Int. J. Mol. Sci. 20: 3945.

Zeng, X., Wang, H., Gong, Z., Huang, J., Pei, W., Wang, X., Zhang, J., and Tang, X. (2015). Antimicrobial and cytotoxic phenolics and phenolic glycosides from *Sargentodoxa cuneata*. Fitoterapia 101: 153–161.

Zhang, J., Subramanian, S., Stacey, G., and Yu, O. (2009). Flavones and flavonols play distinct critical roles during nodulation of *Medicago truncatula* by *Sinorhizobium meliloti*. Plant J. 57: 171–183.

Zhang, J., Subramanian, S., Zhang, Y., and Yu, O. (2007a). Flavone synthases from *Medicago truncatula* are flavanone-2-hydroxylases and are important for nodulation. Plant Physiol. 144: 741–751.

Zhang, X., Xiao, H.-B., Xue, X.-Y., Sun, Y.-G., and Liang, X.-M. (2007b). Simultaneous characterization of isoflavonoids and astragalosides in two *Astragalus* species by high-performance liquid chromatography coupled with atmospheric pressure chemical ionization tandem mass spectrometry. J. Sep. Sci. 30: 2059– 2069.

Zhang, Y., Nie, M., Shi, S., You, Q., Guo, J., and Liu, L. (2014). Integration of magnetic solid phase fishing and off-line two-dimensional high-performance liquid chromatography–diode array detector–mass spectrometry for screening and identification of human serum albumin binders from Radix Astragali. Food Chem. 146: 56–64.

Zhao, X., Mei, W., Gong, M., Zuo, W., Bai, H., and Dai, H. (2011). Antibacterial activity of the flavonoids from *Dalbergia odorifera* on *Ralstonia solanacearum*. Molecules 16: 9775–9782.

Zhou, L., Zhang, Q., Qi, W., Yan, S., Qu, J., Makino, T., and Yuan, D. (2017). Identification of metabolites in human and rat urine after oral administration of Xiao-Qing-Long-Tang granule using ultra high performance liquid chromatography combined with quadrupole time-of-flight mass spectrometry. J. Sep. Sci. 40: 3582–3592.

Zuanazzi, J.A.S., Clergeot, P.H., Quirion, J.-C., Husson, H.-P., Kondorosi, A., and Ratet, P. (1998). Production of*Sinorhizobium meliloti* nod gene activator and repressor flavonoids from *Medicago sativa* roots. Mol. Plant-Microbe Interactions® 11: 784–794.

