## Supplementary Material for "Phenolic compounds in *Medicago truncatula* roots are under the influence of *Agrobacterium fabrum* through its species specific-genes regions"

**Table S1. Primers used in this study**

| *A. fabrum*-specific region | Analysis and region | Forward primer (5' to 3') | Reverse primer (5' to 3') | Reference |
|---|---|---|---|---|
| **Transcriptional fusions** | | | | |
| SpG8-1b | *Atu1416* promoter | | | Meyer et al. 2018 |
| SpG8-2a | *atu3057* promoter | AAGCTTTGACGGTTGTGAACAGCACT | GAGGTGAATACAGGCGGAAA | This study |
| SpG8-2b | *atu3073* promoter | AAGCTTACTGAAACCGACATGAACGC | CCGCCGAGAATTCCGATAGT | This study |
| SpG8-3 | *atu3675* promoter | AAGCTTAAATCGCGTTCTCCAGATGG | GGGTCTCTACTGGCTCGATG | This study |
| SpG8-4 | *atu3817* promoter | TGGTTCCCAGACGTCGTTTA | CAGTTTGATGTAGCGAGCCA | This study |
| SpG8-5 | *Atu3948* promoter | AAGCTTGAAGGTCGGTGGCATTGT | ACGGCTCTTGCTTGTTCG | This study |
| SpG8-7a | *atu4292* promoter | ATCGATGTGCAGAGCTTGCTGACG | CAGGGTGATGTGGAGATCGT | This study |
| SpG8-7b | *atu4299* promoter | AAGCTTCGAGCCATTTCATGAGTGCT | AGGAGACAATGCAACCCGTA | This study |
| | Insertion verification in pGEMT | GTTTTCCCAGTCACGAC | CAGGAAACAGCTATGAC | Promega |
| | Insertion verification in pOT1e | CGGTTTACAAGCATAAAGC | CATTTTTTCTTCCTCCACTAG | Pothier et al. 2007 |
| **Construction of the *A. fabrum*-specific regions deletion mutants (inactivation gene clusters)** | | | | |
| ΔSpG8-1b | Upstream region of *atu1409* | GATGAAGCCGTCAACATCC | AGATCGGTGACGGAGAAG | Lassalle et al. 2011 |
| | Downstream region of *atu1423* | TCCCTCCGGATGAGAACG | GGAGGTGGTCCATGATGT | |
| ΔSpG8-2a | Upstream region of *atu3054* | CTGAATTTTGCGGATGAGC | TATTCGTCGGTTCCTGCG | Lassalle et al. 2011 |
| | Downstream region of *atu3059* | TCCCTCGGAAGTCCGCTT | GTTGCCTCGCTGATGATG | |
| ΔSpG8-2b | Upstream region of *atu3069* | CCGAGTGTGATGATGACGAG | CATGGTGAAGGCAGTTGCTA | This study |
| | Downstream region of *atu3073* | TTCCATAAAGCCCTCATTGG | GTTCGACGAGTTGATGACGA | |
| ΔSpG8-3 | Upstream region of *atu3663* | CCGGTTCTACATCCTGGAAA | GACAACATGCCCCTCTCCTA | Baude et al. 2016 |
| | Downstream region of *atu3693* | CCTGCTCAACAGGCTACTCC | TCTGGAACGTCACCGACATA | |
| ΔSpG8-4 | Upstream region of *atu3808* | ATGACCAGTTGCCAGACCTC | TGATCCGCAGGCCTATATTC | This study |
| | Downstream region of *atu3830* | GTTGACGGCAATATCGTGTG | AGCCAGCAGGATTTTGAGAA | |
| ΔSpG8-5 | Upstream region of *atu3947* | CGTTATCGCTGATCCCATTT | GCTGCCTATTTCCTGTCCAA | This study |
| | Downstream region of *atu3952* | CCTTAACGCAGCTCTTGTCC | CGTGAACCATTTTTGCATTG | |
| ΔSpG8-7a | Upstream region of *atu4285* | GATCGACGAATTGTCGCG | GGCCTGTTGATCTACAGG | This study |
| | Downstream region of *atu4294* | GGGTATCGATAGACGAAG | TCGAATGTCCGAGGATGG | |
| ΔSpG8-7b | Upstream region of *atu4295* | TGCAGCTTGCGATCAATTAC | GCTGAACGAGGTGAAGGAAG | This study |
| | Downstream region of *atu4307* | CAAGCTCCTGACCGACTTTC | GCTGAACGAGGTGAAGGAAG | |

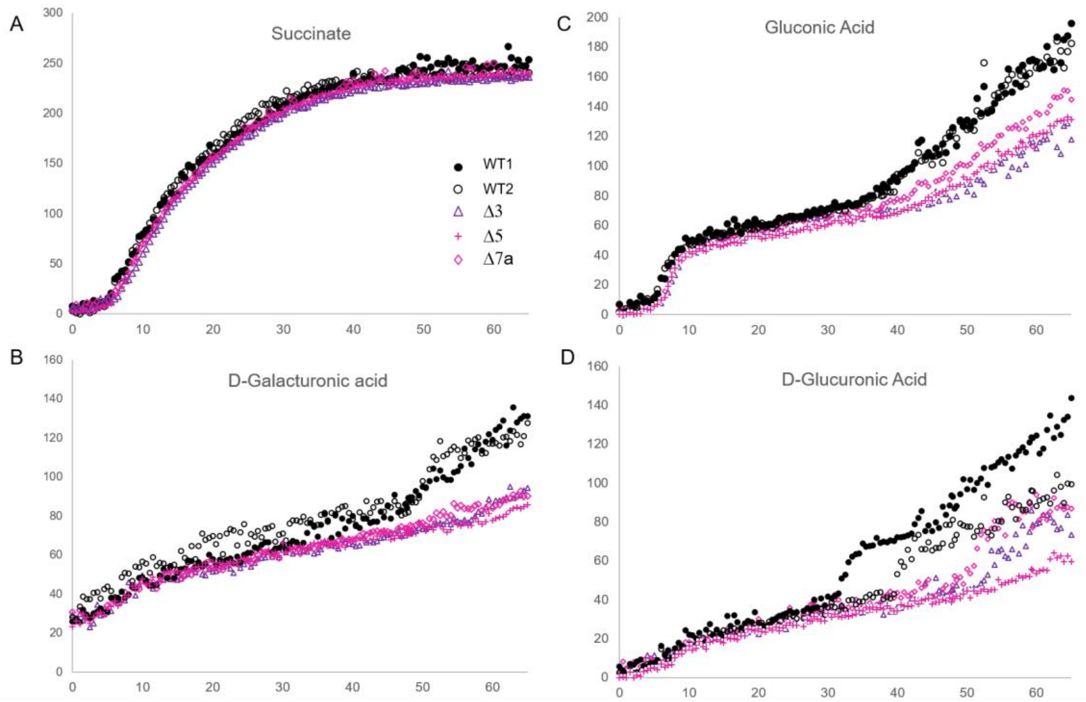


**Fig. S1. Phenotypic changes in species specific mutants compared to wild-type strains.**

Metabolic activity of specific-region deletion mutant strains C58ΔSpG8-3 (=Δ3; △), C58ΔSpG8-5 (=Δ5; **+**) and C58ΔSpG8-7a (=Δ7a; ◇) was compared to wild-type strains WT1 (●) and WT2 (○; see **Table 1** for more information on strains) by using Biolog phenotype microplates. Metabolic activities of the mutant strains are the same as the wild-type strains when grown on succinate (**A**) but differ when grown on hexuronic acids D-galacturonic acid (**B**), gluconic acid (**C**) and D-glucuronic acid (**D**). No difference observed for other carbon sources tested on these microplates (Biolog phenotype MicroArrays PM1 and PM2A).

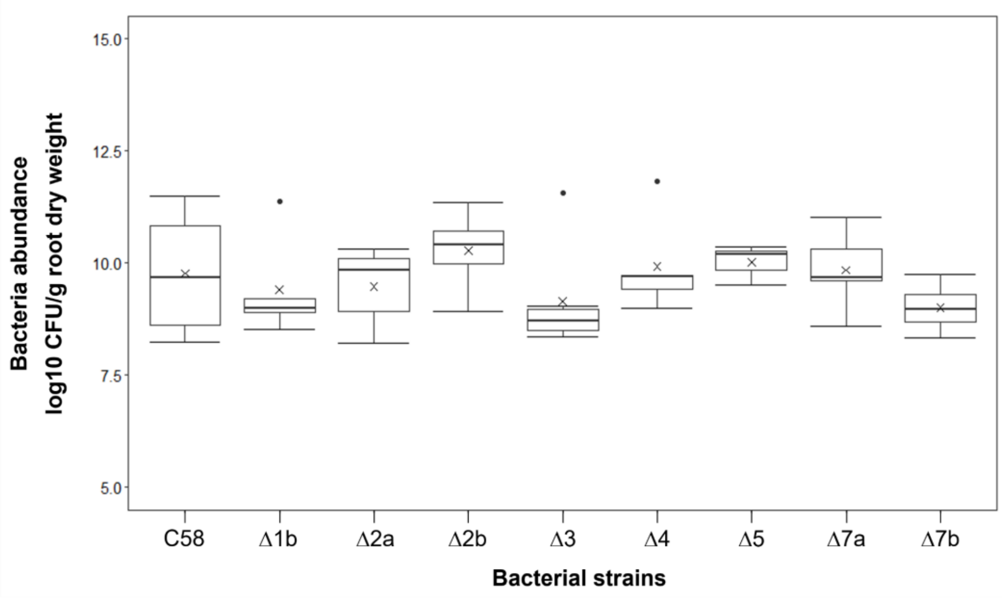


**Fig. S2. Boxplots illustrating *Medicago truncatula* root colonization by *A. fabrum* wild-type and deletion mutant strains.**

For each bacterial strain, the emerging root from seeds of *M. truncatula in vitro* cultivated was inoculated by 5 µL of a suspension at $2.10^5$ bacteria/mL. Bacteria abundance on roots was determined 15 dpi and expressed as log10 cfu (colony-forming units) per gram of root dry weight. Boxes cover 50% of the data. Central lines represent the medians and whiskers represent the minimum and maximum values among non-atypical data. The cross (×) denotes the mean value of the data with at least three replicates. A similar level of root colonization was observed between the wild-type (C58) strain and all the mutant strains (Δ1b = C58ΔSpG8-1b; Δ2a = C58ΔSpG8-2a ; Δ2b = C58ΔSpG8-2b ; Δ3 = C58ΔSpG8-3, Δ4 = C58ΔSpG8-4 ; Δ5= C58ΔSpG8-5 ; Δ7a = C58ΔSpG8-7a ; Δ7b= C58ΔSpG8-7b). No statistically significant differences were observed (Kruskal-Wallis test, $p$ = 0.5026).

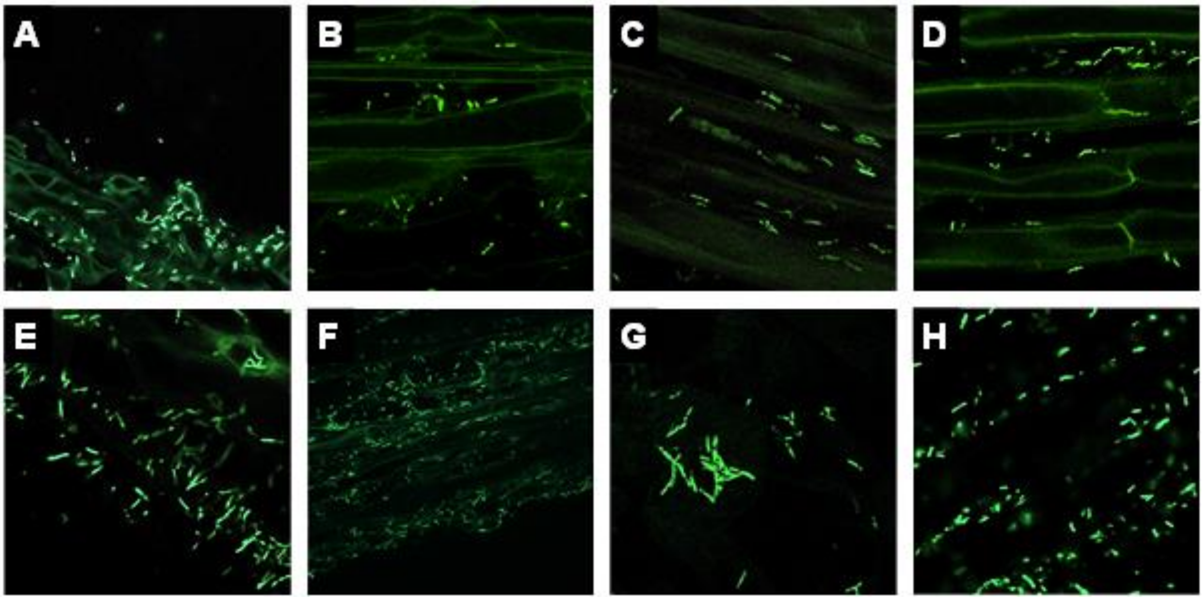


**Fig S3.** ***In planta*** **expression of the** ***A. fabrum*****-specific regions through the expression of their targeted gene on** ***M. truncatula*** **roots. A.**

Expression of SpG8-1b region, represented by expression of *atu1416* gene (*e.g.* transcriptional fusion between *atu1416* promoter region and eGFP gene). **B.** Expression of SpG8-2a region, represented by expression of *atu3057* gene. **C.** Expression of SpG8-2b region, represented by expression of *atu3073* gene. **D.** Expression of SpG8-3 region, represented by expression of *atu3675* gene. **E.** Expression of SpG8-4 region, represented by expression of *atu3817* gene. **F.** Expression of SpG8-5 region, represented by expression of *atu3948* gene. **G.** Expression of SpG8-7a region, represented by expression of *atu4292* gene. **H.** Expression of SpG8-7b region, represented by expression of *atu4299* 7b. Gene expression was monitored by confocal microscopy using transcriptional fusions at 14 dpi. Representative pictures from five plants per specific region are shown. Green fluorescence showed bacteria able to express the corresponding transcriptional fusion. The green autofluorescence of the plants helped to distinguish plant cells. All transcriptional fusions are induced in contact with the entire *M. truncatula* root system.

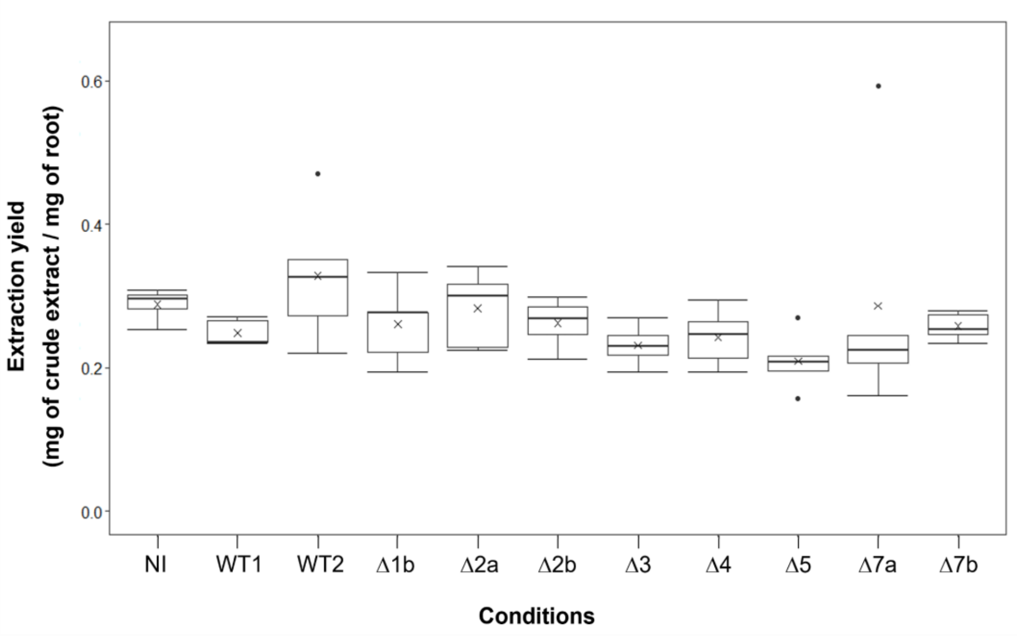


**Fig S4. Boxplots illustrating the extraction yield of metabolites from *Medicago truncatula* root inoculated or not by *A. fabrum* wild-type or deletion mutant strains.**

For each condition 14 dpi, the roots of *M. truncatula in vitro* cultivated were harvested, dried and extracted with methanol. Root crude extracts were weighed and expressed as mg of crude extract per mg of root dry mass. Boxes cover 50% of the data. Central lines represent the medians and whiskers represent the minimum and maximum values among non-atypical data. The cross (x) denotes the mean value of the data (n=4 or 5). Similar amounts of crude extract were obtained from roots, whether *M. truncatula* seedlings were non-inoculated (NI) or inoculated with the wild-type C58 (WT1, WT2) or the deletion mutant strains (Δ1b = C58ΔSpG8-1b; Δ2a = C58ΔSpG8-2a; Δ2b = C58ΔSpG8-2b; Δ3 = C58ΔSpG8-3; Δ4 = C58ΔSpG8-4; Δ5= C58ΔSpG8-5; Δ7a = C58ΔSpG8-7a; Δ7b= C58ΔSpG8-7b). No statistically significant differences were observed (Kruskal-Wallis test, $p = 0.1023$).

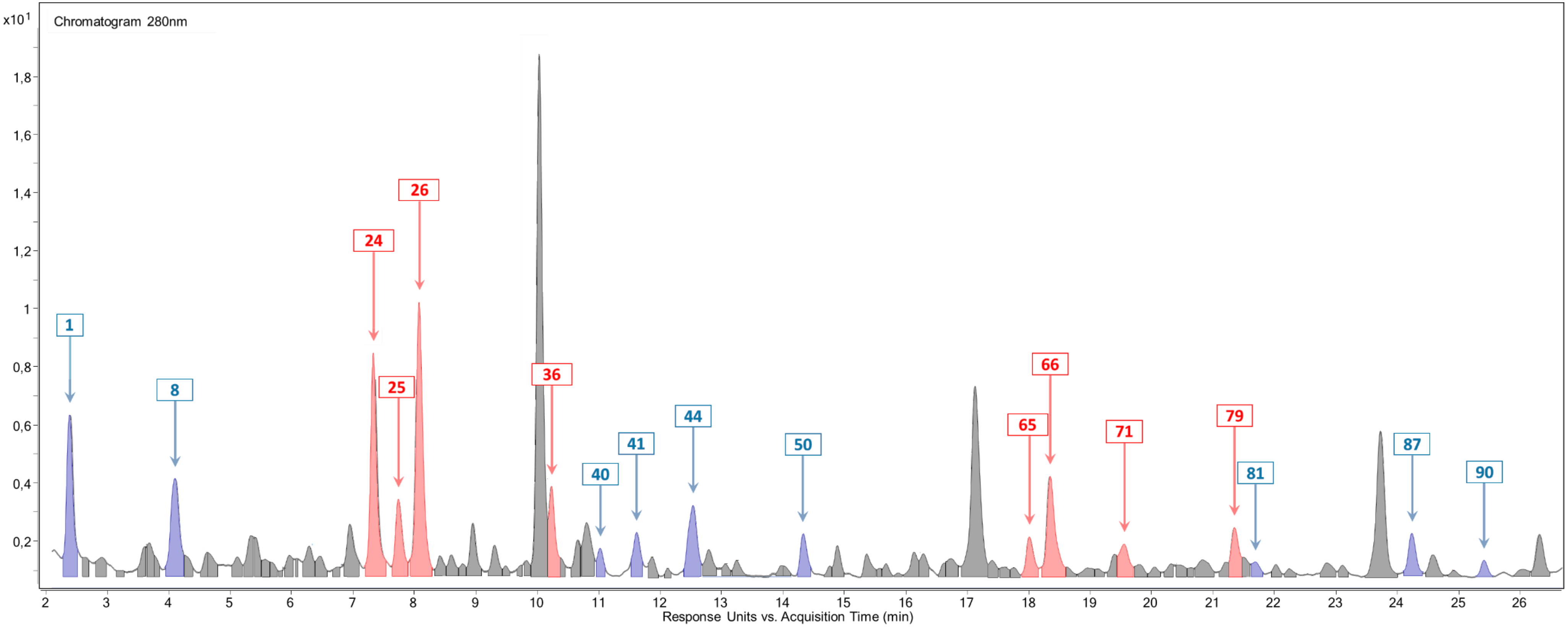


**Fig. S5. A representative UV-chromatogram (280 nm) obtained by UHPLC-UV/DAD-MS QTOF analysis of extracts from *M. truncatula* roots inoculated or not with the wild-type or the deletion mutant strains of *A. fabrum* C58.**

The integrated chromatographic peaks were considered in the metabolite profiling data study (92 peaks). Peak numbering corresponds to metabolite numbering in **Table 3** and in the heatmap in **Fig. 4**. Peaks with an arrow are discriminating compounds with a significant difference ($p<0.05$, Student t-test) between plants inoculated with the wild-type strain (WT condition) compared to the other conditions (the non-inoculated condition and/or the deletion mutants of the *A. fabrum*-specific regions conditions). Peaks in blue correspond to the underabundant compounds, peaks in red correspond to the overabundant compounds compared to the WT condition.

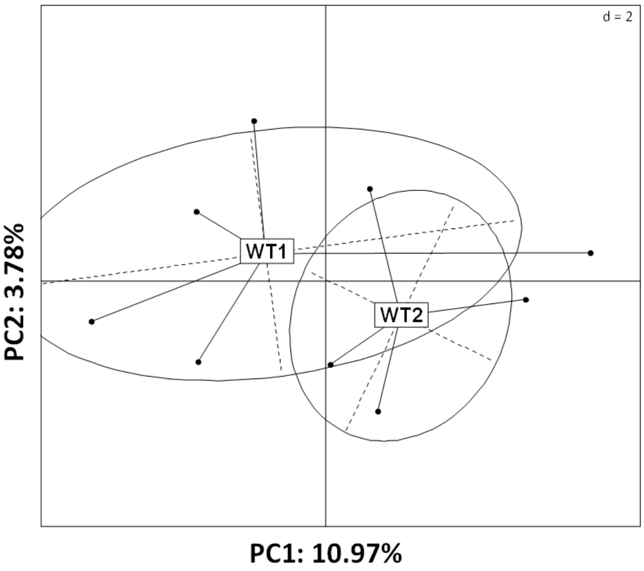


**Fig. S6. PCA score plot of root specialized metabolite profiles of *M. truncatula* seedlings inoculated by the two wild-type strain conditions.**

PCA were performed on the data matrix from chromatographic data at 280 nm obtained from each methanolic extract of *M. truncatula* roots based on peak areas and retention time. Plants were inoculated with *A. fabrum* C58 wild-type strain. WT1: plants inoculated with the wild-type strain, WT2: plants inoculated with the wild-type strain carrying the *ntpII* kanamycin resistance gene. This comparison by PCA did not show any separation of *M. truncatula* root metabolite profiles between these two WT conditions

**SupM1: Identification or annotation of discriminating metabolites**

The detailed interpretations of the spectral data for the structural elucidation of the discriminating compounds are given below.

**Amino acid.** The polar compound **1** eluted at 2.4 min had an ion [M+H]+ at m/z 205.0951 consistent with the molecular formula $C_{11}H_{12}N_2O_2$ and dissociated in MS² mode give ions at m/z 188 [M+H-H2O]+ indicating the loss of a water molecule and at m/z 118 [M+H-87]+ corresponding to an indole moiety according to its accurate mass. Spectral data compared to literature data (Fletcher et al. 2013; Jiang et al. 2011) and to the authentic standard analysis, led to identify this compound as **tryptophan**.

**Flavonoids. Aurones.** Compounds **24** and **26**, eluted respectively at 7.33 and 8.06 min, had identical UV-vis spectra and similar accurate mass at m/z 449.1094 corresponding to [M-H]- ions that both fragmented to give product ions at m/z 287 [M-H-162]-, 269 [M-H-162-18]-, 259 [M-H-162-28]-, indicating the loss of an hexose followed by the loss of -$H_2O$ or –CO respectively. Comparison to literature data (Yoshikawa et al. 1998; Stochmal et al. 2009; Ye et al. 2009; Thuy et al. 2016; Muhammad et al., 2017) led us to propose these compounds as maesopsin glycosides. The analysis of an authentic standard allowed the unequivocal identification of **26** as **maesopsin 4-O-glucoside**. Compound **24** was tentatively identified as an isomer as **maesopsin 6-O-glucoside** (Li et al. 1997; Thuy et al. 2004).

**Flavonoids. Flavanone.** Compound **41** was observed as an ion $[M-H]^-$ at *m/z* 593.1514 yielding in MS² a minor ion at *m/z* 417 $[M-H-176]^-$ indicating the loss of a hexuronic acid and a major ion at *m/z* 255 $[M-H-176 -162]^-$ indicating the loss of a hexose with an accurate mass consistent with the aglycone liquiritigenin. Therefore, combined to literature data (Qiao et al. 2012; Zhou et al. 2017; Ma et al. 2016) and the aglycone standard analysis, this compound was tentatively identified as **liquiritin-7-O-hexuronide**.

**Flavonoids. Flavonol.** Compound **36** was observed as an ion $[M+H]^+$ at *m/z* 771.1950 that yielded fragments at *m/z* 625.1323 $[M+H-146]^+$, 463.0841 $[M+H-146-162]^+$ and 287.0542 $[M+H-146-162-176]^+$ indicating the losses of a deoxyhexose, a [deoxyhexose + hexose] and of a [deoxyhexose + hexose + hexuronic acid], respectively. The negative MS² spectra showed an ion $[M-H]^-$ at *m/z* 769,1841 fragmented in an ion at *m/z* 593,1523 $[M-H-176]^-$ indicating the loss of a hexuronic acid and another ion at *m/z* 285,0404 $[M-H146-162]^-$ indicating a concomitant loss of a hexose and a deoxyhexose. The UV-vis spectrum indicated a kaempferol as aglycone. These data and the literature led to annotate this compound as **kaempferol O-hexuronyl-[deoxyhexosyl]-hexoside** (Budzanowski, 1991), but exact position of the 3 substituents could not be determined.

**Flavonoids. Flavone.** Compound **40** was annotated as a **7,4'-dihydroxyflavone dihexuronide** according to the UV-vis spectrum, fragmentation pattern, literature data (Park et al. 2003; Wong et al. 2009; Ibrahim and Abul-Hajj 1990; Marczak et al. 2016; Singh et al. 2010) and aglycone standard analysis.

Indeed, the fragmentation of the $[M+H]^+$ ion at *m/z* 607.1271 yielded product ions at *m/z* 431 $[M+H-176]^+$ and at *m/z* 255 $[M+H-176-176]^+$ indicating successive losses of two hexuronic acids and leading to the fragment aglycone corresponding to 7,4'-dihydroxyflavone (consistent with the UV-vis spectrum). The $MS^2$ negative spectrum showed a product ion at *m/z* 351.0571 corresponding to the dihexuronide moiety after the loss of the aglycone, leading to the conclusion that the two hexuronic acids were branched together, in accordance with literature data on similar compounds (Marczak et al. 2016). Finally, the hexuronidation on 4'-hydroxyl group of the aglycone was suggested by the hypsochromic shift observed in band I of the UV spectrum (Singh et al. 2010). To our knowledge, this flavone derivative is reported for the first time in the plant kingdom. NMR analysis would be required to definitely characterize the sugar moiety of the compound.

Compound **44** was observed as an ion $[M+H]^+$ at *m/z* 429.0830 giving in $MS^2$ a product ion at *m/z* 253 $[M+H-176]^+$ by losing a hexuronic acid. Spectral data consistent with literature and the aglycone standard analysis led to annotate this compound as **7-4'-dihydroxyflavone 7-O-glucuronide** (Saleh et al. 1982; Singh et al. 2010; Staszków et al. 2011) already described in *Medicago*.

Compound **65** was annotated as **7,4'-dihydroxyflavone-O-[feruloyl-hexuronyl-O-hexuronide]** according to the UV-vis spectrum, fragmentation pattern and aglycone standard analysis. Indeed, the $[M+H]^+$ ion at *m/z* 783.1660 yielded two fragments ions at *m/z* 431 $[M+H-352]^+$ and at *m/z* 353 $[M+H-430]^+$ corresponding to the fragmentation of the molecule in two parts: the aglycone 7,4'-dihydroxyflavone with a hexuronic acid substituent for the first, and a ferulate susbstituted with a hexuronide moiety for the second. A loss of one hexuronic acid was observed from each of these ions, giving product ions at *m/z* 255 $[M+H-352-176]^+$ (the aglycone) and 177 $[M+H-430-176]^+$ (the ferulate). The $MS^2$ negative spectrum showed a product ion at *m/z* 527 $[M-H-254]^-$ corresponding to the loss of the aglycone leading to the conclusion that the two hexuronides and ferulate moiety were all linked together (Guy et al. 2009; Arni et al. 2010). The other product ion at *m/z* 333 $[M-H-254-194]^-$ (loss of aglycone and ferulate) demonstrated that the two hexuronides were connected to each other. To our knowledge, this type of flavone derivative was reported for the first time in the plant kingdom. NMR analysis would be required to definitely characterize the sugar moiety of the compound.

Compound **50** was observed as an ion $[M+H]^+$ at *m/z* 623.1209, consistent with the molecular formula $C_{27}H_{26}O_{17}$. $MS^2$ positive spectrum showed daughter ions at *m/z* 447 $[M-H-176]^+$ and 271 $[M-H-176-176]^+$ indicating successive losses of hexuronide moieties. These data and the UV-vis spectrum indicated an apigenin as aglycon. Furthermore, in $MS^2$ negative spectrum, the formation of a product ion at *m/z* 351 resulting from the loss of 270amu (the aglycone apigenin) showed that the two hexuronides are attached to each other. A hypsochromic shift was observed in band I of the UV-vis spectrum compared to that of the apigenin, which seemed to show that the substituent is on the nucleus B. Thus, the glucuronidation must be on the 4'-hydroxyl group of apigenin (Mabry et al. 1970). Indeed, the apigenin 7-O-diglucuronide already described in *Medicago truncatula* (Jasiński et al. 2009) was found in our extracts at another retention time of 13.24 min (corresponding to compound **47**). Its

UV-vis spectrum, however, showed an $\lambda_{max}$ at 266 and at 340 nm, indicating a substitution occurring in ring A of the aglycone, unlike compound **50** (Stochmal et al. 2001a). Therefore, this data combined to literature data and the aglycone standard analysis, led to tentatively identify this compound as **apigenin 4'-O-dihexuronide**. This compound is described for the first time in the plant kingdom. As before, NMR analysis is needed to confirm the structure and precise the configuration of this new compound.

Compound **66** was observed as an ion $[M\text{-}H]^-$ at *m/z* 475.0905 consistent with the molecular formula $C_{22}H_{20}O_{12}$. $MS^2$ spectrum showed fragment ions at *m/z* 299 $[M\text{-}H\text{-}176]^-$ and at *m/z* 284 $[M\text{-}H\text{-}176\text{-}15]^-$ indicating the loss of an hexuronic acid followed by a loss of $-CH_3$ from the aglycone, a diosmetin according to UV-vis spectrum. Data obtained compared to literature data (Beninger and Hall 2005; Li et al. 2016b; Ferreres et al. 2014; Greenham et al. 2003; Yu et al. 2015a) and the aglycone standard analysis led us to annotated this compound as a **diosmetin-O-hexuronide**. However, exact position of the substituent (on 3'- or 7-hydroxyl group) could not be determined.

Compound **79** showed a UV spectrum leading to suspect an apigenin aglycone susbtituted with an HCA compound. In MS, it was observed as a protonated ion at *m/z* 799.1615 $[M+H]^+$, which was further fragmented in $MS^2$ producing a series of daughter ions at *m/z* 447 $[M+H\text{-}352]^+$ (loss of hexuronic acid and ferulate), *m/z* 353 $[M+H\text{-}446]^+$ (loss of hexuronic acid and apigenin), *m/z* 271 $[M+H\text{-}352\text{-}176]^+$ (ion of apigenin aglycone) and *m/z* 177 $[M+H\text{-}446\text{-}176]^+$ (ion of ferulate moiety). Exactly the same fragmentation pattern was described in the literature for a compound detected in leaf extract of *Medicago truncatula.* This compound was annotated as **apigenin O-[feruloyl-glucuronopyranosyl-O-glucuronopyranoside]** (Jasiński et al. 2009).

**Flavonoids. Isoflavone.**

Compound **71** was observed as an ion $[M+H]^+$ at *m/z* 431.1348 consistent with the molecular formula $C_{22}H_{22}O_9$. It yielded in positive $MS^2$ a product ion at *m/z* 269 $[M+H\text{-}162]^+$ which revealed the loss of an hexose from an aglycone which could be a formononetin according to the $\lambda_{max}$ of the UV-vis spectrum (230, 252 and 296 nm). The negative $MS^2$ showed a deprotonated ion at *m/z* 475 $[M\text{-}H+HCOO]^-$ that fragmented to give ions at *m/z* 267 $[M\text{-}H\text{-}162]^-$ corresponding to a loss of a hexose and the formic acid and at *m/z* 252 $[M\text{-}H\text{-}162\text{-}15]^-$ indicating the loss of the $-CH_3$ from the aglycone formononetin. The analysis of the standard compound ononin confirmed the identification of this **formononetin-7-O-glucoside**, besides being already described in *M. truncatula* cell suspension cultures (Farag et al. 2007).

Compound **87** showed in the positive MS spectrum an ion$[M+H]^+$ at *m/z* 547.1517 consistent with the molecular formula $C_{26}H_{26}O_{13}$. $MS^2$ spectrum revealed product ions at *m/z* 299 $[M\text{-}H\text{-}162\text{-}86]^+$ and 284 $[M\text{-}H\text{-}162\text{-}86\text{-}15]^+$corresponding to a loss of a glucoside malonate and the loss of the $-CH_3$ from the aglycone afrormosin, consistent with the molecular formula and UV-vis spectrum. All these data compared to literature led us to annotated this compound as **afrormosin-7-O-glucoside-6'-O-**

**malonate**, already described in *M. truncatula* (Tibe et al. 2011; Zhang et al. 2014; Farag et al. 2007, 2008; Zhang et al. 2007b)
